# Overlapping attentional networks yield divergent behavioral predictions across tasks: Neuromarkers for diffuse and focused attention?

**DOI:** 10.1101/713339

**Authors:** Esther X.W. Wu, Gwenisha J. Liaw, Rui Zhe Goh, Tiffany T.Y. Chia, Alisia M.J. Chee, Takashi Obana, Monica D. Rosenberg, B.T. Thomas Yeo, Christopher L. Asplund

## Abstract

Attention is a critical cognitive function, allowing humans to select, enhance, and sustain focus on information of behavioral relevance. Attention contains dissociable neural and psychological components. Nevertheless, some brain networks support multiple attentional functions. Connectome-based Predictive Models (CPM), which associate individual differences in task performance with functional connectivity patterns, provide a compelling example. A sustained attention network model (saCPM) successfully predicted performance for selective attention, inhibitory control, and reading recall tasks. Here we constructed a visual attentional blink (VAB) model (vabCPM), comparing its performance predictions and network edges associated with successful and unsuccessful behavior to the saCPM’s. In the VAB, attention devoted to a target often causes a subsequent item to be missed. Although frequently attributed to attentional limitations, VAB deficits may attenuate when participants are distracted or deploy attention diffusely. Participants (n=73; 24 males) underwent fMRI while performing the VAB task and while resting. Outside the scanner, they completed other cognitive tasks over several days. A vabCPM constructed from these data successfully predicted VAB performance. Strikingly, the network edges that predicted better VAB performance (positive edges) predicted *worse* selective and sustained attention performance, and vice versa. Predictions from the saCPM mirrored these results, with the network’s negative edges predicting better VAB performance. Furthermore, the vabCPM’s positive edges significantly overlapped with the saCPM’s negative edges, and vice versa. We conclude that these partially overlapping networks each have general attentional functions. They may indicate an individual’s propensity to diffusely deploy attention, predicting better performance for some tasks and worse for others.

**Significance statement:** A longstanding question in psychology and neuroscience is whether we have general capacities or domain-specific ones. For such general capacities, what is the common function? Here we addressed these questions using the attentional blink (AB) task and neuroimaging. Individuals searched for two items in a stream of distracting items; the second item was often missed when it closely followed the first. How often the second item was missed varied across individuals, which was reflected in attention networks. Curiously, the networks’ pattern of function that was good for the AB was bad for other tasks, and vice versa. We propose that these networks may represent not a general attentional ability, but rather the tendency to attend in a less focused manner.

## Introduction

Attention is a critical cognitive function, allowing humans to select, enhance, and sustain focus on information of behavioral relevance. Visual attention plays numerous roles in different contexts, and it has been fractionated both behaviorally and neurally (Chun, Golomb, & Turk-Browne, 2011; Desimone & Duncan, 1995; Egeth & Yantis, 1997). In addition to such separable components, however, some brain networks support attentional processing across multiple domains (Asplund, Todd, Snyder, & Marois, 2010; Corbetta & Shulman, 2002; Duncan, 2010; Tamber-Rosenau, Dux, Tombu, Asplund, & Marois, 2013; Yeo, Krienen, et al., 2015). Recent studies using Connectome-based Predictive Models (CPM) support this idea. In a CPM approach, individual differences in behavioral performance are accounted for as a function of whole-brain functional connectivity patterns, after which performance for novel individuals can be predicted from fMRI data (Shen et al., 2017). Such predictions also apply across tasks. A sustained attention network model (saCPM) (Rosenberg, Finn, et al., 2016) could predict individual differences in performance for selective attention (Rosenberg, Hsu, Scheinost, Constable, & Chun, 2018), inhibitory control (Fountain-Zaragoza, Samimy, Rosenberg, & Prakash, 2019), and reading recall (Jangraw et al., 2018).

Here we constructed a CPM for the visual attentional blink (VAB), aiming to test whether that model could predict performance on a variety of tasks and to compare its predictions and network features to the saCPM’s. In a VAB paradigm, participants search for two items in a stream of distractors; they often fail to perceive the second item, but only when it closely follows the first (200-500 ms) (Raymond, Shapiro, & Arnell, 1992). The VAB is critically dependent on attention, as no deficit occurs when the first item is ignored. Individuals differ in their VAB severity (rate of second item detection failures), and these differences are typically large and stable (Dale, Dux, & Arnell, 2013). It is unclear, however, which cognitive and neural factors underlie them. Numerous theoretical explanations have been proposed for the VAB, including a temporary loss of control (Di Lollo, Kawahara, Shahab Ghorashi, & Enns, 2005) or bottleneck-like processing limitations (Chun & Potter, 1995). VAB magnitude also correlates only weakly with most other attention tasks (Skogsberg et al., 2015).

Intriguingly, VAB performance sometimes improves when attention to its primary detection task is reduced. Such reductions can be due to manipulation (Olivers & Nieuwenhuis, 2005, 2006) or dispositional factors (Dale & Arnell, 2010, 2015; Thomson, Ralph, Besner, & Smilek, 2015), and are thought to cause more diffuse attentional deployment. In particular, mind-wandering is associated with better VAB performance, though it has the opposite association for many other attention tasks (Gonçalves et al., 2017; Hu, He, & Xu, 2012; Robertson, Manly, Andrade, Baddeley, & Yiend, 1997; Smilek, Carriere, & Cheyne, 2010).

The VAB likely involves many factors (Dux & Marois, 2009), but which are reflected in individual differences of brain network function? To address this question using a CPM approach, we scanned 73 individuals while they performed the VAB task and while they rested. Resting state data allowed us to assess whether any predictive functional architecture persisted when participants were not engaged in attention-demanding tasks (Finn et al., 2015; Rosenberg, Finn, et al., 2016; Yoo et al., 2017). Outside the scanner, the same individuals completed cognitive tasks assessing sustained attention, selective attention, and fluid intelligence. We constructed a visual attentional blink CPM (vabCPM), from which we could make and assess predictions about the tasks. If attentional capacity predicts VAB performance, we would expect *positive* associations between vabCPM predictions for behavior and observed performance in other attention tasks. Conversely, if diffuse attentional tendencies predict VAB performance, we might find significant *negative* associations between predicted and observed behavior for other attention tasks. For external validation, we made and assessed predictions about our tasks, including the VAB, using a sustained attention CPM (Rosenberg, Finn, et al., 2016). We then investigated and compared the networks associated with each model to better understand their relationship and potential psychological functions.

## Materials and Methods

The present study included numerous tasks, with a primary focus on the visual attentional blink (VAB). Additional tasks provided critical context and points of comparison for understanding the individual differences in VAB performance and neural features. These additional tasks included those related to goal-directed attention and fluid intelligence. Many of these tasks are conceptually linked to the VAB (Dux & Marois, 2009), and many have also been studied themselves using Connectome-based Predictive Modeling (Finn et al., 2015; Rosenberg, Finn, et al., 2016; Rosenberg et al., 2018). To facilitate comparisons across these different tasks, we re-coded all behavioral performance measures such that positive numbers indicated better performance (e.g. higher accuracy or faster reaction times; see details below).

## Experimental design

### Participants

Eighty-two participants with self-reported normal or corrected-to-normal vision and normal hearing were recruited from the National University of Singapore (NUS) community. These individuals began a six-session study that included numerous behavioral and neuroimaging components, a subset of which are reported and analyzed here. Eight participants did not continue with the experiment after the first practice session (Session 0) and were excluded from the following analyses. One participant who did not achieve a target discrimination score of 75% in the main VAB task (Session 1) was also excluded. Thus, unless otherwise stated, the following analyses included data from 73 participants (24 males) between the ages of 19-30 (M = 22.25, SD = 1.84). All participants provided written informed consent in accordance with a protocol approved by the NUS Institutional Review Board and received monetary compensation.

### Stimulus presentation

All sessions took place either inside the MR scanner or in the laboratory over a span of 3 weeks (see Table 1-1 of Extended Data for complete and detailed experimental protocol). Inside the scanner, stimuli were presented at a viewing distance of 91 cm on a 32-inch LCD monitor (NordicNeuroLab, Bergen, Norway) with a screen refresh rate of 60 Hz, connected to a MacBook Air (OS 10.12.1) running PyschoPy (Peirce, 2007). Participants made responses using an MR-compatible button box. In the laboratory, stimuli were presented at a distance of 57 cm on a 22-inch LCD monitor (Samsung SyncMaster 2233) with a screen refresh rate of 60 Hz using an NVIDIA Quadro FX 3450/4000 SD graphics card on Windows 7. Participants’ responses were captured on a standard computer keyboard. Auditory stimuli were presented using PyschoPy (Peirce, 2007) binaurally through Creative headphones.

**Table 1.**
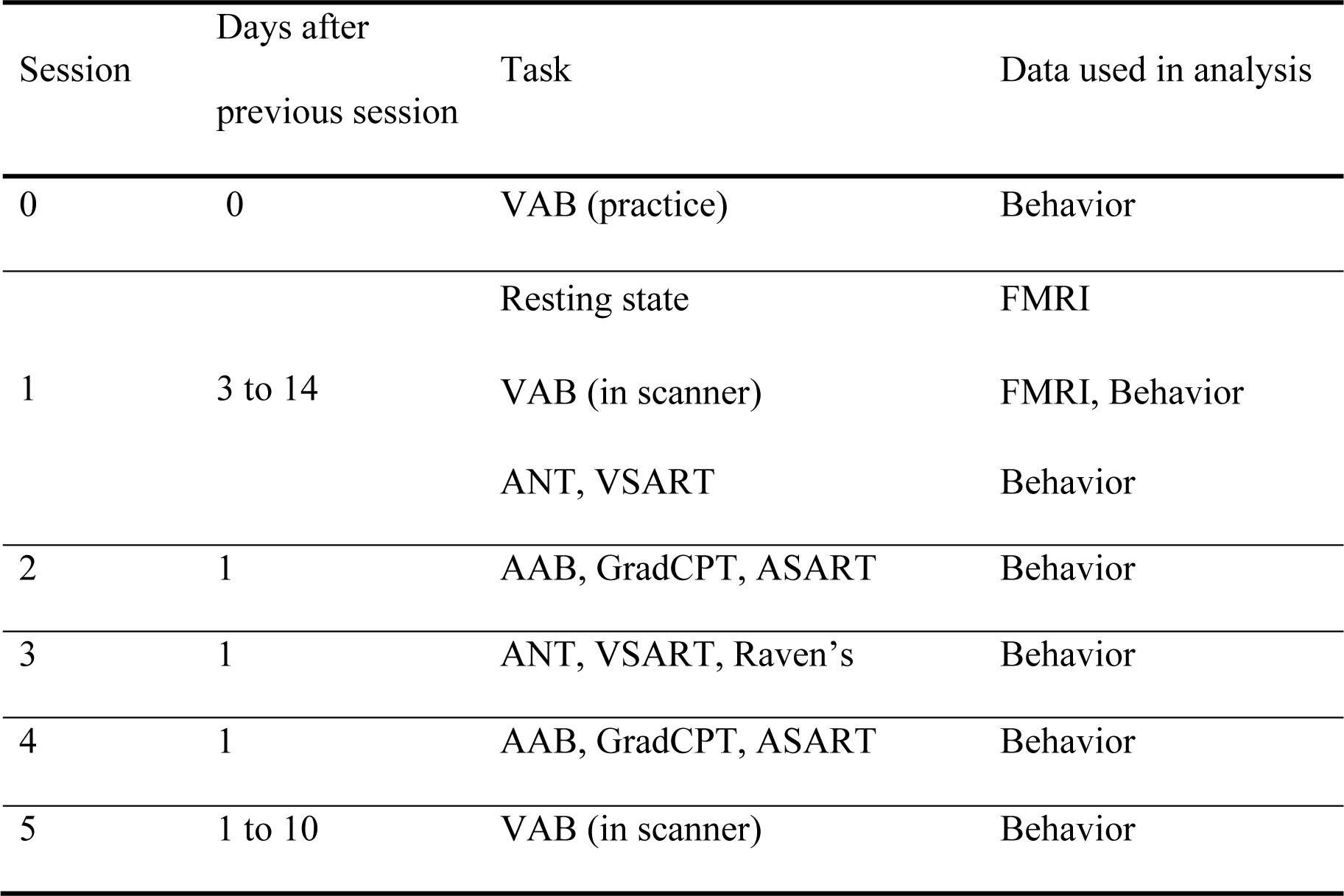
Schedule of tasks and data used in analysis. Participants were encouraged to take breaks between the tasks to prevent fatigue. With the exception of the Raven’s Progressive Matrices test, each task was performed twice on non-successive days. Task domains and tasks were as follows. Attentional Blink: Visual Attentional Blink (VAB) and Auditory Attentional Blink (AAB). Sustained Attention: Visual Sustained Attention to Response Task (VSART), Auditory Sustained Attention to Response Task (ASART), and Gradual-onset Continuous Performance Task (GradCPT). Selective Attention: Attentional Network Task (ANT). Fluid Intelligence: Raven’s Progressive Matrices test (Raven’s).

### Overview of task domains and specific tasks

The tasks in this study all investigated cognitive processing, primarily different forms of attention (Table 1). Each task is detailed below, organized by task domain. The task domains included the Attentional Blink, Sustained Attention, Selective Attention, and Fluid Intelligence.

### Attentional Blink

Participants completed a visual attentional blink (VAB) and an auditory attentional blink (AAB) task. They were designed to be generally similar (Figure 1). Both versions were built around target discrimination and a probe detection, with each item embedded within a stream of distractors. Our main task of interest is the VAB, for which neuroimaging data was collected concurrently. For the AAB and all other tasks, only behavioral data were collected.

**Figure 1.**
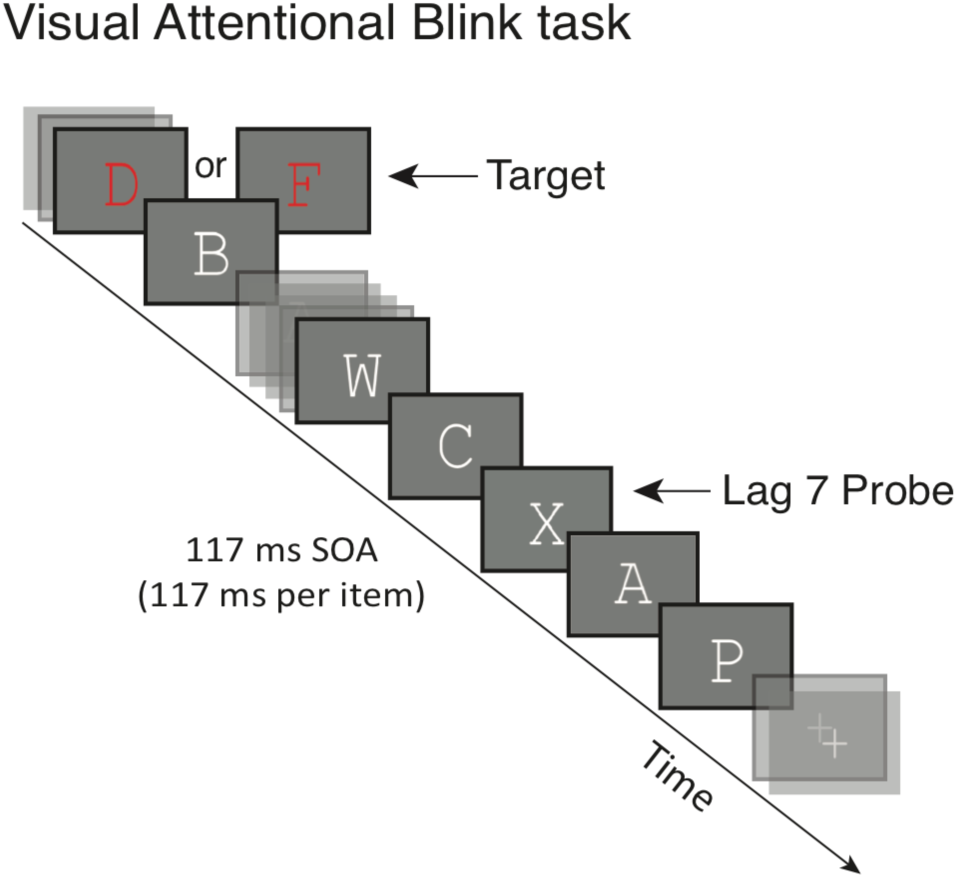
Experimental paradigm for the visual attentional blink (VAB) task. Participants identified a target and then detected a probe within a stream of distractors, responding when prompted at the conclusion of the stream. The target was a red letter, the probe was a white letter X, and distractors were other white letters. The auditory attentional blink (AAB) task was similar, save targets were complex tones, the probe was a high-pitched pure tone, and distractors were other pure tones. SOA = stimulus onset asynchrony.

For the VAB, stimuli consisted of upper-case letters presented in Courier New Font on a dark gray background (Figure 1). Targets were red letters D or F, whereas the probe was a white letter X. Distractors were white letters save D, F, X, I, L, O, and Q. Targets and probes were embedded within a rapid serial presentation stream, with no items repeated during each trial. The stimulus onset asynchrony (SOA) between successive items in the 16-stimulus stream was 117 ms (no gap). A target appeared during every trial at serial position 3, 4, or 5, whereas the probe appeared during 75% of trials. When present, the probe appeared at lags 1, 2, 3, 5, 7, or 9 relative to the target, with the same number of trials per lag condition in each block. The first three lags were expected to be within the AB window, whereas the last three were expected to be outside it. In a small percentage of trials (14%), an irrelevant surprise stimulus (randomly selected from a set of 24 grayscale male faces and 24 colorful objects) was presented at lags 2 or 6 relative to the target. These surprise stimuli are not relevant to this current study; trials containing surprises were excluded from the following analyses and are not reported further.

Each 6.25-second trial began with the presentation of a white fixation cross (0.8° x 0.8°) for 500 ms, which became larger (1.0° x 1.0°) and turned yellow to signify that the rapid serial visual presentation (RSVP) stream would begin in 750 ms. Participants searched this RSVP stream (1867 ms) for the target and probe, which they then indicated by button box press after a blank gap (233 ms) at the trial’s conclusion. A maximum of 2.9 s was given for participants to respond to both the target and probe response prompts. After this period, no further responses were recorded and the white fixation cross returned until the start of the next trial. Failure to respond was rare: No session had more than 0.16% no-target response trials or 0.64% no-probe response trials averaged across participants, and no participant had more than 3.57% (target) or 4.17% (probe) no-response trials in any given session. The timing between each trial was optimized for functional Magnetic Resonance Imaging (fMRI). As such, it followed an exponential distribution with a range of 1.25-10 s and mean of 3.75 s. Each session contained six blocks of 28 trials each, with the trials presented in a pseudorandom order. The 168 trials took approximately 40 minutes to complete, including breaks. Before the main experimental blocks in each session, each participant completed 3 practice blocks of 8 trials each. The first block contained targets but no probes; the second block contained probes but no targets; and the third block contained both.

For the AAB, targets were low-pitched or high-pitched complex tones comprised of five log-related frequencies (794 to 1260 Hz, or 1349 to 2142 Hz), whereas the probe was a 4000 Hz pure tone. Distractors were 19 pure tones of log-related frequencies ranging from 697 to 2911 Hz. Sound stimuli were adjusted to have equal mean absolute amplitudes, after which the probe and distractor intensities were set to 45% and 30% of the target intensity (∼70 dB). These values were based on performance in Obana, Lim, & Asplund (under review) and additional pilot tests. As in the visual task, a small percentage of auditory trials (14%) contained an irrelevant surprise stimulus, randomly selected from a set of 24 sounds (including an alarm, a cough, and spoken letters), presented at lags 2 or 6 relative to the target. Again, surprise trials were not analyzed here. With the exception of a change in stimuli, trial structure was identical to the VAB.

Each 6.25-second trial began with a reminder of the target and probe sounds, which were played for 110 ms each with a gap of 85 ms. After 750 ms, the rapid auditory stream (RAP) began, through which participants searched for the target and probe. Similar to the visual task, the identity of the target and the presence of the probe were indicated by keypress after a blank gap (233 ms). Response prompts for the target and probe then appeared, with a maximum allowed time of 2.9 s for both responses. The timing between each trial was fixed at 0.75 s.

Except for the blank gap and the response prompts, a white fixation cross (0.8° x 0.8°) was shown on the screen throughout the block. Each session contained three blocks of 56 trials each, with the trials presented in a pseudorandom order. The 168 trials took approximately 25 minutes to complete, including breaks. Before the main experimental blocks in each session, each participant completed 3 practice blocks of 8 trials each. The first block contained targets but no probes; the second block contained probes but no targets; and the third block contained both.

Before beginning each practice and task block of the auditory task, participants could play the target and probe sounds as many times as desired.

### Sustained attention

To better understand VAB performance in relation to other forms of attention and their neural underpinnings, we also ran three sustained attention tasks. These paradigms included the visual and auditory versions of the Sustained Attention to Response Task (SART; Robertson, Manly, Andrade, Baddeley, & Yiend, 1997), which we adapted from Seli, Cheyne, Barton, & Smilek (2012), and the Gradual-Onset Continuous Performance Task (GradCPT; Esterman, Noonan, Rosenberg, & DeGutis, 2013; Rosenberg, Noonan, DeGutis, & Esterman, 2013). The SART has been frequently used to examine moment-to-moment fluctuations of sustained attention, requiring participants to make continuous responses to most stimuli but withhold responses to a few. However, due to its trial-based structure, which may provide a short ‘break’ between trials and not tax attention sufficiently, the GradCPT was later designed to present images that gradually transition from one to the next using a linear pixel-by-pixel interpolation. The GradCPT has been shown to show reliable and large interindividual variability amongst high-functioning young adults, such as those in our sample (Rosenberg, Finn, et al., 2016; Rosenberg et al., 2013).

In the VSART, participants were presented single digits, one after another, in the center of the display screen. They were asked to press the spacebar if they saw a number from 1 to 9, but withhold their response if they saw the number 3. Each digit was presented for 250 ms, followed by an encircled “x” mask for 900 ms. Digits were presented in Symbol font in white, against a black background, at sizes 0.57°, 1.03°, 1.43°, 1.89°, 2.35° of visual angle. The order of the digits and their sizes were randomized. Participants completed 675 trials (∼ 13.5 mins) in each session.

For the ASART, stimuli consisted of spoken single numbers. As in the visual task, participants were asked to press the spacebar when they heard the numbers 1 to 9, but to withhold their response if they heard the number 3. The numbers were presented in random order, each for 250 ms, following by a pink noise mask of 900 ms. Throughout the experiment, participants maintained fixation on a cross in the middle of the display. Each session consisted of 675 trials (∼13.5 mins).

The GradCPT (Rosenberg et al., 2013) consisted of images that gradually transitioned from one to the next using a linear pixel-by-pixel interpolation (ISI = 800 ms). Images consisted of 10 mountain and 10 city scenes, randomly presented with 10% and 90% probability, respectively, without repeats in consecutive images. Participants were instructed to press the spacebar when a city scene was presented, but to withhold their response when a mountain scene was presented. To tax sustained attention sufficiently, the GradCPT was performed in a single block over a relatively long duration (15 min).

### Selective attention

To understand the AB’s relationship to other selective attention tasks and their neural underpinnings, we also employed the Attentional Network Task (ANT), by Fan, McCandliss, Sommer, Raz, & Posner (2002). This paradigm was designed to test three separable components of selective attention: alerting, orienting and executive control (Posner & Petersen, 1990) within a single experimental session. Stimuli consisted of 5 black lines (some with arrowheads) arranged horizontally in a row, against a grey background. The target, always an arrow in the center, was flanked on each side by (i) two arrows pointing in the same direction as the target (congruent condition), (ii) two arrows pointing in the opposite direction from the target (incongruent condition), or (iii) two black lines without arrowheads (neutral condition). Each line or arrow measured 0.55° horizontally, and the space between two adjacent objects measured 0.06°. To trigger attention orienting, all stimuli were presented either 1.06° above or below a central fixation cross. Participants were asked to keep their eyes fixated on the fixation cross and respond whether the target was pointing right or left.

Each trial started with a central fixation cross (400 – 1600 ms), followed by a warning cue (100 ms), a second central fixation cross (400 ms), and finally, the stimuli consisting of target and flankers presented either above or below a central fixation cross. Target and flankers were presented for 1700 ms, or until a response was made, whichever was shorter. A fixation cross was then presented until the end of trial (4000 ms after the first fixation period). For the warning cue, four types of cues were presented: (i) no cue (a central fixation cross, similar to that presented during the first and second fixation period, was presented), (ii) a center cue (an asterisk was presented in the center, thus alerting the participant to the impending stimuli presentation), (iii) double cue (two asterisks were presented above and below a central fixation cross, at both possible locations of the target), and (iv) a spatial cue (an asterisk was presented at the impending location of the target). Participants completed 3 blocks of 96 trials (4 cue conditions x 2 target locations x 2 target directions x 2 repetitions), with trials presented in random order. The entire task lasted about 30 min.

### Fluid intelligence

As a final comparison domain for understanding VAB performance and the associated neural underpinnings, we measured fluid intelligence. Participants completed a shortened, nine-item version (Form A; Bilker et al., 2012) of the original 60-item Raven’s Standard Progressive Matrices (Raven, Raven, & Court, 1998). The task was completed on a laboratory computer, and there were no response time limits. The task consists of pattern matching questions designed to measure abstract reasoning skills, and it has been typically used in clinical settings as a non-verbal test of fluid intelligence.

## Statistical analyses for behavioral data

Participants had to meet a minimum target discrimination (letter D or F?) score of 75% for the VAB and 60% for the AAB to be included in the final sample. As a result, 73 sets of data were available for all tasks, with the exception of the AAB task (n=71). For the AAB, two additional participants were excluded because their target discrimination performance did not meet the minimum threshold in each session. For all behavioral comparisons, *p*-values were based on two-tailed comparisons. We also did not correct any behavioral comparisons for multiple corrections, as we wanted to find any normality violations and used the task correlations to identify any relationships that might affect our CPM results.

The computation of each behavioral measure is detailed in the following sections, again organized by task domain. To obtain stable behavioral metrics, we computed a ‘best score’ for each task metric. When data was available across two different sessions, the final score was averaged across both sessions. (When data was available only from a single session, the final score set to that session’s.) We assessed whether the distribution of ‘best scores’ for each metric departed from normality using Jarque-Bera tests. As many normality violations were found, we used Spearman correlations of the ‘best scores’ to compare each pair of tasks. For tasks with two sessions, we also calculated test-retest reliability. Due to the aforementioned normality violations, Spearman correlations were again used.

### Attentional blink

The VAB and AAB deficits were calculated in the same way. For each participant, we first computed the mean probe detection accuracy for each lag condition, contingent upon correct identification of the target. Probe detection accuracy scores were then averaged across short-lag (lags 1, 2, or 3) and long-lag (lags 5, 7, 9) conditions. The former condition was expected to be within the attentional blink window, whereas the latter condition was expected to be outside it. The AB deficit was computed by regressing out the long-lag scores from the short-lag scores of each participant (short-lag scores ∼ long-lag scores), and then saving the residuals (MacLean & Arnell, 2012). Larger and positive values indicated smaller attentional deficits, and thus better task performance. Note that simply subtracting the short-lag from the long-lag scores yielded highly similar VAB deficit scores (*r*(71) = .955). Visual AB scores (VABresid) were obtained from sessions 1 and 5, and the auditory AB scores (AABresid) were obtained from sessions 2 and 4.

### Sustained attention

D-prime values were computed for the VSART, ASART and GradCPT, with larger values indicative of better performance in sustained attention. VSART scores (VSARTdprime) were obtained from sessions 1 and 3, ASART scores (ASARTdprime) were obtained from sessions 2 and 4, and GradCPT scores (GradCPTdprime) were obtained from sessions 2 and 4.

### Selective attention

To measure overall task performance in the ANT task, we computed the mean error rate (ANTerr) across all trials and the intra-individual variability of RTs (ANTrtvar) for each participant. Intra-individual RT variability was computed as the standard deviation divided by mean of correct-trial RTs. Arguably, this metric is a more sensitive measure of task performance than mean error rate (Rosenberg et al., 2018; Wojtowicz, Berrigan, & Fisk, 2012), with higher RT variability being linked to lower accuracy in ANT tasks (Adolfsdottir, Sorensen, & Lundervold, 2008; Lundervold et al., 2011).

We also calculated metrics for each attentional network in the ANT, which putatively reflect their efficiencies. To do so, we compared the RTs between different trial conditions for each participant (Fan et al., 2002; Rosenberg et al., 2018). For the alerting network, efficiency was computed by subtracting the mean RT of double-cue condition from the mean RT of the no-cue condition (ANTaert). Larger values would indicate faster responses due to the cue and thus better task performance. For the orienting network, efficiency was computed by subtracting the mean RT of spatial-cue condition from the mean RT of the center-cue condition (ANToert), thus larger positive values would also imply faster responses due to the cue and better task performance. For the executive control network, efficiency was computed by subtracting the mean RT of the congruent condition from the mean RT of the incongruent condition (ANTcert). In this case, smaller values would imply less interference by the flanker arrows and better task performance. For all computations of efficiency with RTs, only trials with correct responses were included.

For easier comparisons with the other behavioral measures, we re-coded the raw values of mean error (ANTerr), RT variability (ANTrtvar), and executive control efficiency (ANTcert), multiplying them by -1 such that larger values would also imply better task performance. ANT scores were obtained from sessions 1 and 3.

### Fluid intelligence

Accuracy scores for the nine-item Raven’s test (RavensAcc) were computed for each participant, with higher scores implying better performance.

**Table 2.**
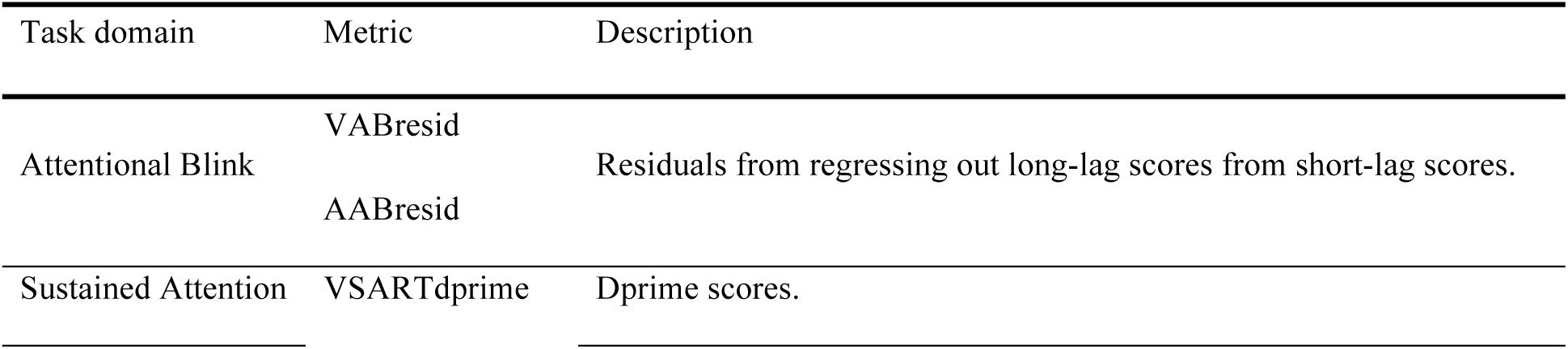
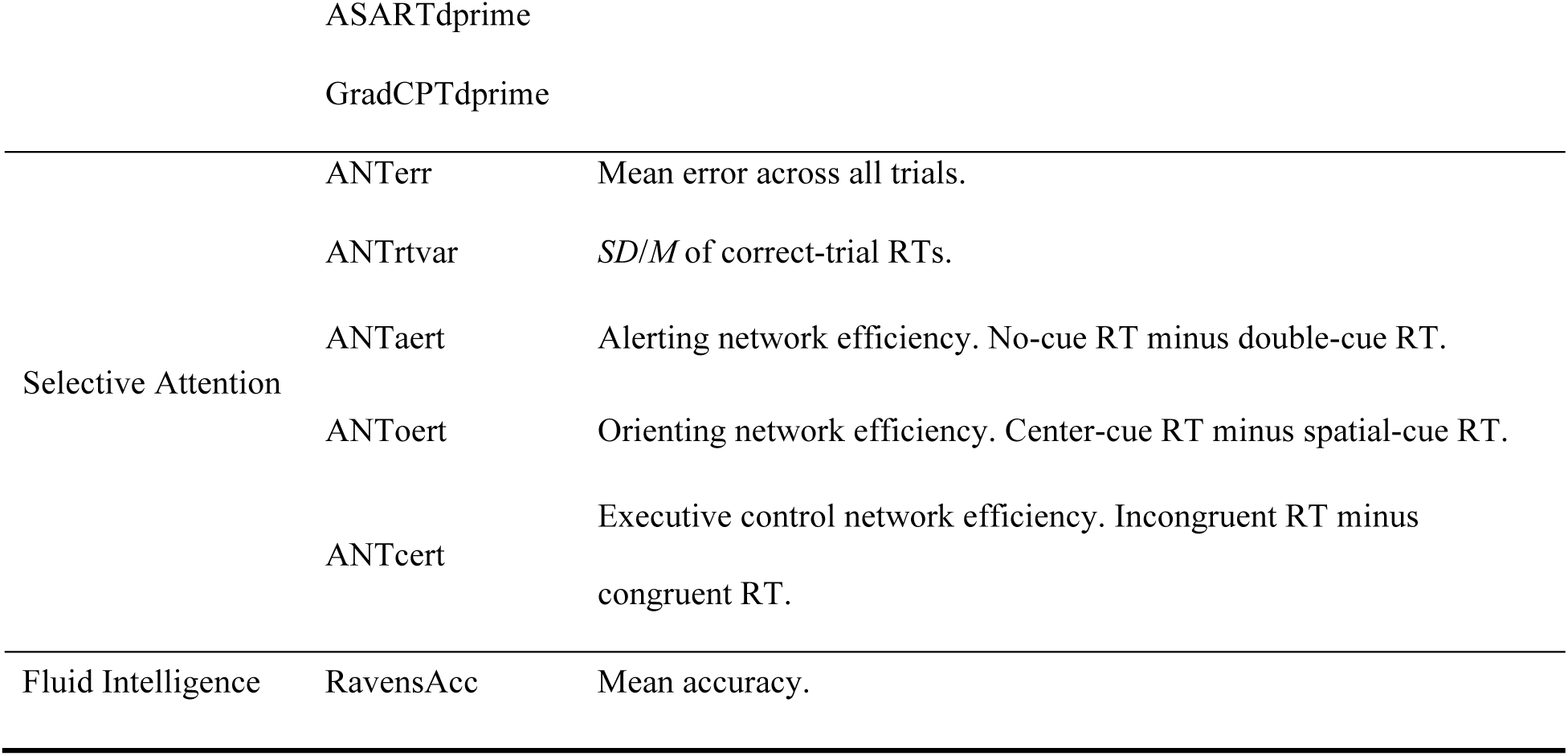
Summary of behavioral metrics, including their task domain and a description of their calculation. Raw values for ANTerr, ANTrtvar and ANTcert were re-coded such that larger values indicated better task performance.

## MRI data collection and initial processing

### Acquisition

MRI data were acquired at the Clinical Imaging Research Centre (Singapore) on a Siemens 3T MAGNETOM Prisma MRI scanner (Siemens, Erlangen, Germany) with a 32-channel head coil. Scanning parameters were adapted from the Human Connectome Project (HCP), and were chosen to ensure that full-brain coverage, including the cerebellum, was achieved for each participant. MRI data were acquired in Sessions 1 and 5 of the study, with the first session’s data analyzed here. Each fMRI session started with a 5-min anatomical localizer scan, followed by four 8-min resting-state scans, six 5.25-min task-based (VAB task) scans, and a 5-min anatomical scan. During the resting-state runs, participants were asked to maintain fixation at a cross displayed in the center of the screen.

### Imaging parameters

A 3D high-resolution (1 mm x 1 mm x 1 mm) T1-weighted MPRAGE pulse sequence was used to obtain whole-brain anatomical images for each participant, allowing for subsequent normalization to standard space. For each participant, 128 1-mm thick contiguous sagittal slices (0.5 mm skip; 1 x 1 mm in-plane resolution) were acquired. Other scanning parameters included a repetition time (TR) of 2300 ms, an effective echo time (TE) of 2.22 ms, a flip angle of 8° and 260 mm field of view.

Functional MRI data were acquired with a multiband echoplanar imaging (MB-EPI; CMRR release R2015; Feinberg et al., 2010; Moeller et al., 2010; Xu et al., 2013) sequence with a MB acceleration factor of 8. 768 whole-brain images were obtained for each resting-state run while 504 images were acquired for each task-based run. T2*-weighted images were acquired using a TR of 625 ms, a TE of 33.2 ms and FA of 50°. Interleaved slices (imaging matrix = 64 x 64) were collected using a 220 mm field of view, with slice thickness at 2.50 mm (no gap). The effective voxel size was thus 2.5 x 2.5 x 2.5 mm^3^.

### Preprocessing

Both task and localizer data were pre-processed using a previously published pipeline for functional connectivity analyses (Fong et al., 2019; Kong et al., 2019) publicly available at https://github.com/ThomasYeoLab/CBIG/tree/master/stable_projects/preprocessing/CBIG_fMRI_Preproc2016. Pre-processing steps across resting-state and task-based runs were the same, except when mentioned otherwise. First, the initial four frames from each run were removed to aid with BOLD signal stabilisation. Motion correction using FSL’s MCFLIRT was then applied such that runs with more than 50% of the frames exceeding a motion threshold were discarded to ameliorate any contributions of head motion. The motion threshold was defined as frame displacement (FD) > 75 and frame-to-frame intensity (DVARS) > 0.2 (Power, Barnes, Snyder, Schlaggar, & Petersen, 2012). From this step, one task-based run (and no resting-state runs) was removed from our data. FSL’s bbregister function was used for intrasubject registration of the T1 anatomical images to the T2*-weighted images. The best run for each subject was used as the registration file across all functional runs. Subsequently, motion parameters and their derivatives, the global whole brain signal, the white matter signal, the cerebral spinal fluid signal and linear trends were regressed out. Additionally, for task-based runs, we regressed out the haemodynamic response signal aligned to trial onset times. Frames with excessive motion, identified earlier on, were interpolated over (Power et al., 2014), and a temporal filter was applied to retain frequencies between 0.009 and 0.08 Hz. The resulting BOLD signal was projected to fsaverage6 surface space and spatially smoothed with an isotropic Gaussian kernel of 6 mm (FWHM).

### Functional connectivity

Functional connectivity was evaluated in fsaverage6 surface space for 400 cortical regions (Schaefer et al., 2017) and in MNI152 volumetric space for 19 subcortical regions (including the brain stem, and the left and right hemispheres of the accumbens area, amygdala, caudate, cerebellum, ventral diencephalon, hippocampus, pallidum, putamen, and thalamus), with a total of 419 parcellations. For each run, the mean time course of all the parcellations were correlated using Pearson’s product moment correlation, resulting in a 419 (rows) x 419 (columns) correlation matrix, with (419 x 419 – 419) / 2 = 87,571 unique values. Each cell in the correlation matrix represents a functional connection (edge) between a pair of parcellations. Fisher r-to-z transformation was applied to increase normality (Van Dijk et al., 2010). For each participant, Fisher-transformed matrices for all four resting-state scans and all six task-based scans were averaged separately, forming two functional connectivity (FC) matrices: resting-state FC (RSFC) and Visual Attentional Blink task FC (VABFC).

### Motion control

To reduce the effects of motion on functional connectivity, we adapted motion control procedures from Rosenberg et al. (2018) to remove edges that were correlated with motion. For each participant, we measured the following five motion parameters from their resting-state and task-based scans: (i) maximum displacement, (ii) maximum rotation, (iii) mean frame-to-frame displacement, (iv) mean frame displacement (FD), and (v) mean frame-to-frame intensity (DVARS). Spearman’s rank correlation was computed across participants between each edge in the FC matrix and each motion parameter. To be comparable with Rosenberg et al. (2018), in which 72.7% of edges remained after controlling for motion, we removed edges where *r* > .275 (two-tailed *p* < .02), leaving 69,617 edges, or 79.50% of the initial 87,571 edges. Edges were removed from both VABFC and RSFC if they met the criteria for removal in either FC matrix.

## General approach for Connectome-based Predictive Models (CPM)

We adapted the Connectome-based Predictive Model (CPM) approach (Finn et al., 2015; Rosenberg, Finn, et al., 2016; Rosenberg et al., 2018; Shen et al., 2017) to predict individual differences in behavior from functional connectivity information (Figure 2). In the following section, we first describe our CPM procedure and then summarize our different models. Data from all 73 participants were included in these analyses (71 for models involving the AAB score, AABresid).

**Figure 2.**
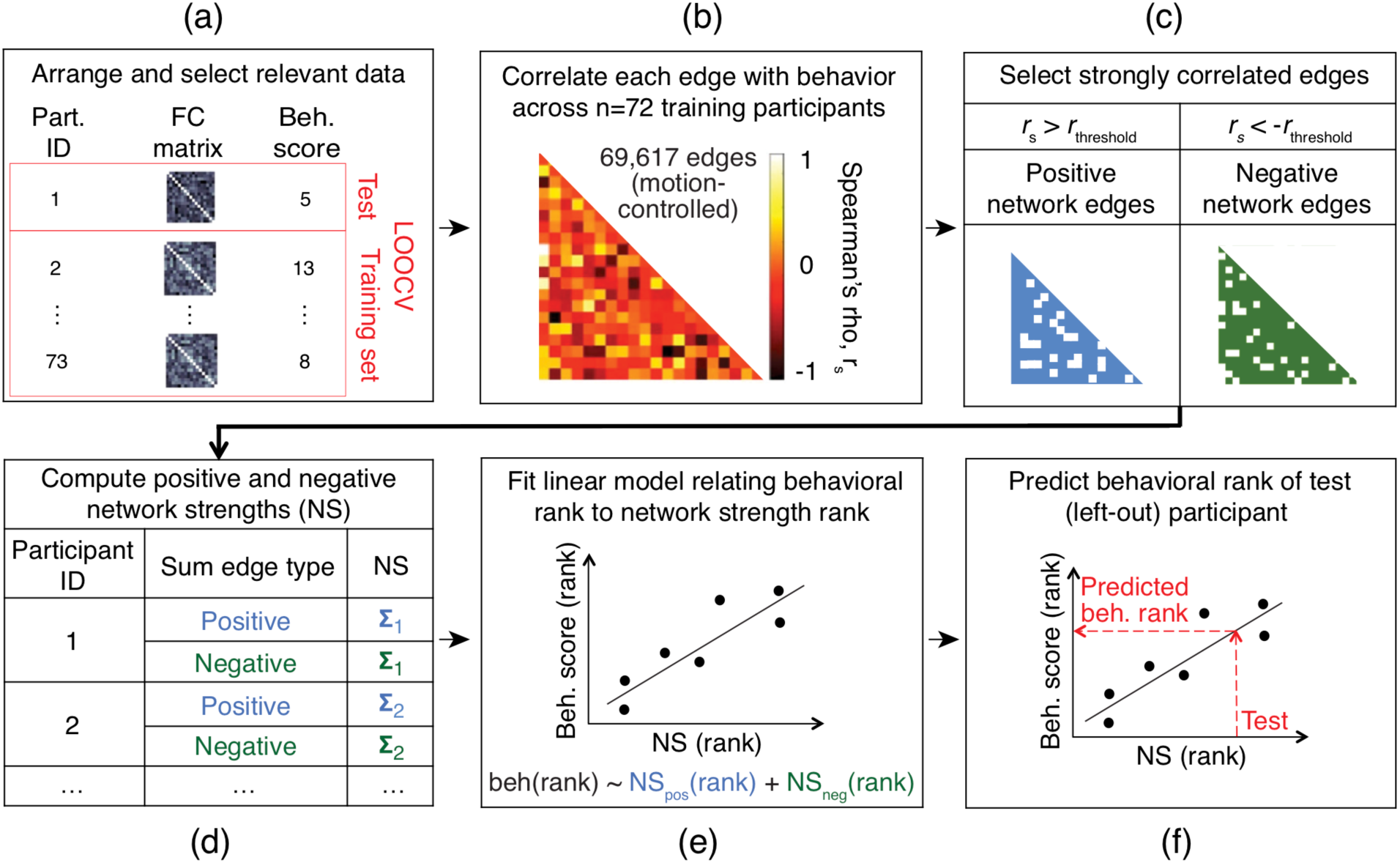
Procedure for Connectome-based Predictive Model (CPM) construction in the current study (adapted from Shen et al., 2017). CPMs predict individual differences in behavior from functional connectivity information. (a) Functional connectivity matrices and a behavioral score of interest for each participant were calculated. One pair was held out of model construction for each round of leave-one-out cross-validation (LOOCV) (Webb et al., 2011). (b) Functional connectivity edges were correlated with behavior across participants. (c) Edges that correlated most strongly, either positively or negatively, were selected. (d) Values from selected edges were summed separately for positive and negative network edges, yielding two network strengths for each participant. (e) A linear regression model relating (rank) network strengths to (rank) behavioral scores was computed. (f) The model was tested on a novel, out-of-sample participant (the individual left out in step (a)). After repeating steps a-f for each participant, the model was evaluated by correlating the predicted behavioral scores with the actual scores.

### Model training

We first selected an FC matrix (i.e., VABFC or RSFC) and a behavioral score (e.g., VABresid). As part of the leave-one-out cross-validation procedure (Figure 2a), we then set aside the data for one participant as test data and proceeded to train the model on the remaining data (*n* = 72). To identify edges that most strongly correlated with behavior, we computed Spearman’s rank correlations between each unique edge in the FC matrix and the behavioral score across 72 participants (Figure 2b), yielding 69,617 Spearman’s rho values (*r*_s_). Edges positively related to behavior (positive network edges) were identified as those whose *r*_s_ was greater than a pre-defined threshold, *r*_threshold_, and edges negatively related to behavior (negative network edges) were defined as those whose *r*_s_ was less than -*r*_threshold_ (Figure 2c).

Next, we computed network strengths (NS) for each participant by summing up values in their individual FC matrices across all positively and negatively correlated edges (Figure 2d), resulting in 72 sets of positive NS and negative NS values. Subsequently, we converted the positive NS, negative NS, and behavioral scores to rank space by ordering them according to participants’ values. Finally, we formulated a multiple linear regression model (Figure 2e) with positive NS rank and negative NS rank as independent variables and behavioral rank as the dependent variable i.e., Behavior (rank) ∼ NS_positive_ (rank) + NS_negative_ (rank).

### Model test

We proceeded to predict the behavioral score (in rank space) of the test participant (Figure 2f) by applying the training model to his/her FC matrix. Using the same positive and negatively correlated edges identified from training, positive NS and negative NS were first computed by summing up values in the FC matrix across the respective edges. Following this step, positive and negative NS values of the test participant were ranked against the relevant NS values of the training participants, and entered into the multiple linear regression model to predict a behavioral rank.

### Model evaluation

As part of the leave-one-out cross-validation procedure, the above training and prediction steps were repeated on all participants (*N* = 73) such that each participant was left out of training once, resulting in 73 sets of predicted behavioral ranks. To measure the predictive power of the model, we obtained correlations between the predicted and observed values, controlling for motion (Rosenberg et al., 2018). To this end, partial Spearman’s rank correlation was computed between predicted and observed behavioral ranks, with the motion parameters (see section on Motion control) included as covariates. Where FC matrices were different for training and test, motion parameters from both the resting-state and task-based scans were included (i.e., ten covariates); where the same FC matrix was used, only motion parameters from the relevant scan were included (i.e., five covariates). *P*-values for model evaluation were left uncorrected, as most of the comparisons represented planned replications of previous work, and the pattern of predictions across tasks was more informative than single model predictions. For other neuroimaging-based statistical tests, we corrected for multiple comparisons, as detailed in the respective Results sections (Functional connectivity and Network overlap).

## CPM model specification for data from the current study

For each of the eleven behavioral scores (e.g., VABresid, AABresid, VSARTdprime, etc.), we repeated the CPM procedure to train and test four different types of models. Two model types (vabCPM model variants) were trained using FC information and behavioral responses acquired from the main VAB task (i.e., VABFC and VABresid), and used to predict behavioral performance of novel participants using either their VAB *task* (VABFC; Model type A), or their *rest* (RSFC; Model type B) FC information. With these models, we sought to identify attentional networks that were predictive of good VAB performance (positive network) and poor VAB performance (negative network), and then to determine whether these networks generalized to make similar predictions about other tasks. Two other model types (task-specific model variants) were trained using task-specific behavioral data and either VAB *task* (VABFC; Model type C) or *rest* (RSFC; Model type D) information. Behavioral performance was then predicted from the same type of FC data that was used during training. With these models, we tested whether predictive networks for each task could be predicted from FC data that was unrelated to that task; any such predictive networks would be useful for identifying and comparing the edges that are predictive of performance on a given task.

### Model type A (train on VABFC and behavioral data, predict with VABFC data)

For training, the functional connectivity matrix VABFC and behavioral score VABresid were used to select edges and form the linear model. For test, the training model was applied to the VABFC matrix of the left-out participant. For evaluation, the predicted behavior from the leave-one-out procedure was correlated with the behavioral score from a selected task.

### Model type B (train on VABFC and behavioral data, predict with RSFC data)

As in the previous model, VABFC and VABresid were used for training, but for test, the training model was applied to the RSFC matrix of the left-out participant. For evaluation, the predicted behavior was correlated with the behavioral score from a selected task.

### Model type C (train and test on VABFC data)

During training, VABFC and the behavioral score from a selected task were used to form the model. During test, the training model was applied to VABFC of the left-out participant. For evaluation, the predicted behavior was correlated with the observed score of the selected task.

### Model type D (train and test on RSFC data)

For training, the functional connectivity matrix RSFC and the behavioral score from a selected task were used to form the model. For test, the training model was applied to the RSFC of the left-out participant. During evaluation, the predicted behavior was correlated with the observed score of the selected task.

### Controls

For the primary analyses in the study, we implemented the CPM procedure using edge selection cutoffs (*r*_threshold_ = .232, *p* = .05) previously used in Rosenberg et al. (2018). As an exploratory control, we also investigated whether our predictions were reasonably stable across edge selection cutoff values. To do so, we repeated the leave-one-out cross-validation procedure with *r*_threshold_ ranging from .005 to .5, in steps of .005. Thus, in total, the CPM procedure was repeated (11 behavior x 4 models x 101 edge selection thresholds) 4,444 times.

As *p*-values from LOOCV procedures can be biased, we verified the significance of our models using permutation testing (Shen et al., 2017). A null distribution with 1000 iterations was generated for each *r*_threshold_. For each iteration, we randomly shuffled participants’ behavioral scores and repeated the prediction steps above. *P*-values (uncorrected for this exploratory analysis) were computed as the proportion of permutation *r*_s_ with values greater than the observed *r*_s_.

## Sustained Attention CPM from Rosenberg et al. (2016)

To compare our predictions and networks with another attention-related model, we applied the Sustained Attention CPM (saCPM) (Rosenberg, Finn, et al., 2016) to our FC data. The saCPM was constructed using FC data computed with 268 parcellations (Shen, Tokoglu, Papademetris, & Constable, 2013). As our FC data was computed with 419 parcellations (Schaefer et al., 2017), we transformed the Shen parcellations to Schaefer parcellations in MNI152 space (91x109x91, 2mm voxels). For each Shen parcellation, we located the corresponding Schaefer parcellation at a corresponding spatial location, excluding those that accounted for less than 10% of the voxels in the Shen parcellation. Next, we mapped the saCPM edges in the following way: if Shen parcellation A mapped to Schaefer parcellations 1, 2, and 3 and Shen parcellation C mapped to Schaefer parcellations 7 and 8, a functional connection (edge) between Shen parcellations A and C would be mapped to Schaefer edges 1-7, 2-7, 3-7, 1-8, 2-8, and 3-8. As before, edges that were removed previously due to motion were also removed in the mapped saCPM edges.

We computed network strengths by taking the dot product between the saCPM edges and our FC matrix (VABFC or RSFC), and entered the result into a linear model: Behavior ∼ NS_positive_ + NS_negative_. Note that these were motion-controlled FCs (69,617 edges), but during evaluation, we did not implement partial correlation with motion parameters as co-variates, following Rosenberg, Finn, et al. (2016). As the saCPM model was trained using FC data and dprime scores acquired during the GradCPT task, predicted scores from the model were also dprime scores. We evaluated the result of applying the saCPM to our FC data by computing Spearman’s rank correlation between the predicted scores and the scores from each of our behavioral tasks.

## Network overlaps and edge locations

### Degree of overlap

To better understand the relationships across CPMs, we calculated the percentage of network overlap for model pairs. First, for each model, we identified edges that were common across all iterations of the leave-one-out procedure; we reasoned that these edges were most representative of the model. For each pair of models, we expressed the number of edges in common percentages of the number of edges in each model, and we then computed the average of the two percentages. These percentages were calculated separately for positive-positive, negative-negative, positive-negative, and negative-positive network overlaps.

To statistically assess whether the edges for each pair of models significantly overlapped, we used the hypergeometric cumulative density function to determine the probability of drawing up to *x* out of *K* possible items with *n* drawings without replacement, from a population of size *M* (Rosenberg, Zhang, et al., 2016). The procedure was implemented with the hygecdf function in (*MATLAB*, 2014), as *p* = 1 – hygecdf(*x*, *M*, *K*, *n*), with x as the number of overlapping edges, M as the total number of edges, *K* as the number of edges from one model, and *n* as the number of edges from the other model. To control for multiple comparisons, FDR correction was applied across the complete set of tests (Benjamini & Hochberg, 1995).

### Anatomical location of networks and overlaps

To determine the anatomical locations of network edges and the overlaps between the vabCPM and saCPM networks, we first grouped the 419 parcellations into network groups. The parcellations were matched to 17 network labels (Yeo et al., 2011), from which they were aggregated into eight cortical groups (Yeo, Tandi, & Chee, 2015) and a subcortical group. For each pair of network groups (e.g. Visual and Salience / Ventral Attention), we computed the number of connections (network edges) between them. We then reported this value as a percentage of the total number of possible connections between that pair of network groups.

## Results

In the sections below, we first report the behavioral results and the functional connectivity matrices. We then present the behavioral predictions based on models derived from the present dataset, followed by the behavioral predictions based on an external model (the saCPM; Rosenberg et al., 2016). After comparing the pattern of predictions, we compare the degree of overlap across the predictive networks. Finally, we examine the anatomical locations of the overlaps between the vabCPM and saCPM networks.

## Behavioral performance

We found a robust visual attentional blink (VAB) in each session, with substantially impaired probe detection performance for the shorter lags (1, 2, 3) but not the longer lags (5, 7, 9) (Figure 3). For subsequent individual differences analyses, the VAB deficit was defined as short-lag performance when controlling for long-lag performance (see Methods), a definition the results supported. An AAB was also evidenced, though it was both smaller and less robust across sessions.

**Figure 3.**
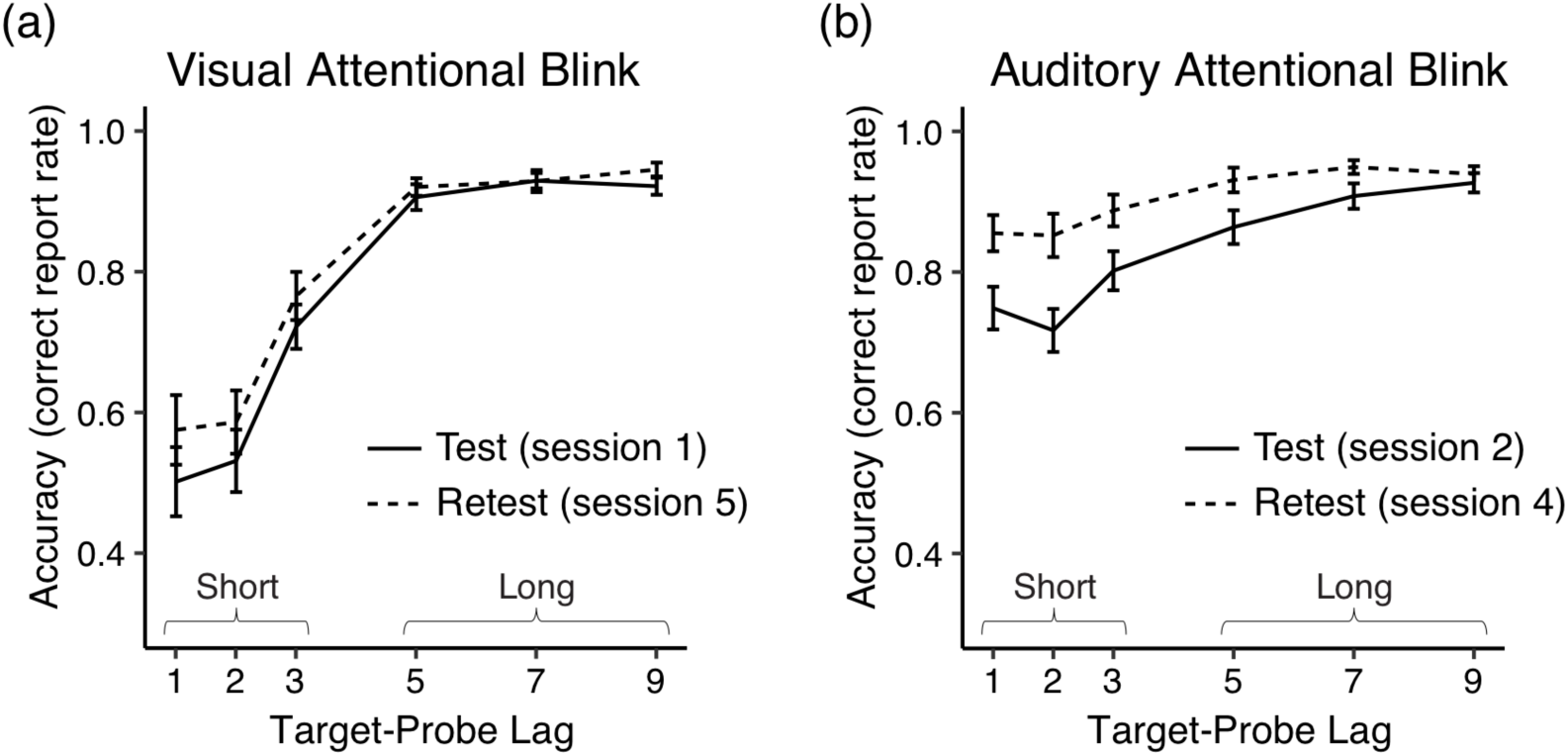
Accuracy scores (probe hit rates) for the VAB and AAB. Note the substantial impairment for the short target-probe lags (1, 2, and 3) in each session, especially for the VAB. Error bars represent standard error of the mean (SEM).

Test-retest reliability was high for most metrics, and their ranges were reasonable (Table 3). Jarque-Bera tests of normality revealed that several behavioral scores were not normally-distributed (see relevant plots in Figure 4). Hence we adopted non-parametric approaches in our subsequent analyses and models.

**Figure 4.**
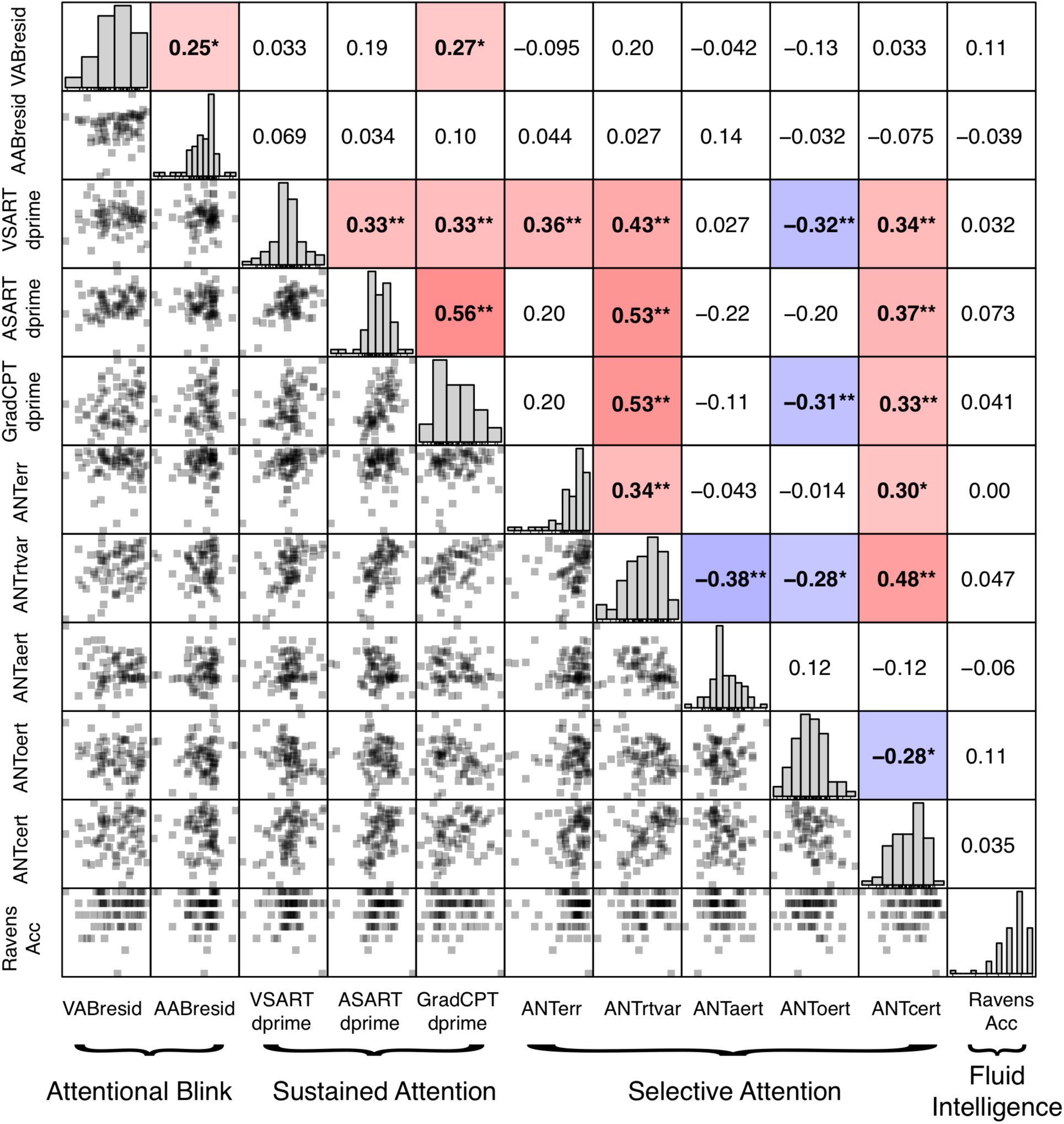
Behavioral score distributions and correlations. (Above diagonal) Spearman correlation coefficients for pairs of behavioral metrics. Most significant correlations were found within task domains, but some metrics correlated across the Sustained and Selective Attention domains. In contrast, Attentional Blink and Fluid Intelligence metrics largely did not correlate with other metrics. Since the purpose of this analysis was to identify any behavioral relationships that might explain subsequent CPM results, no correction for multiple comparisons was applied. \**p* < .05, \*\**p* < .01. Red and blue shading indicates positive and negative relationships, respectively. (Diagonal) Histograms of behavioral data. The behavioral data had been re-coded so that larger values indicate better task performance for each measure. (Below diagonal) Scatterplots for each pair of behavioral metrics.

**Table 3.**
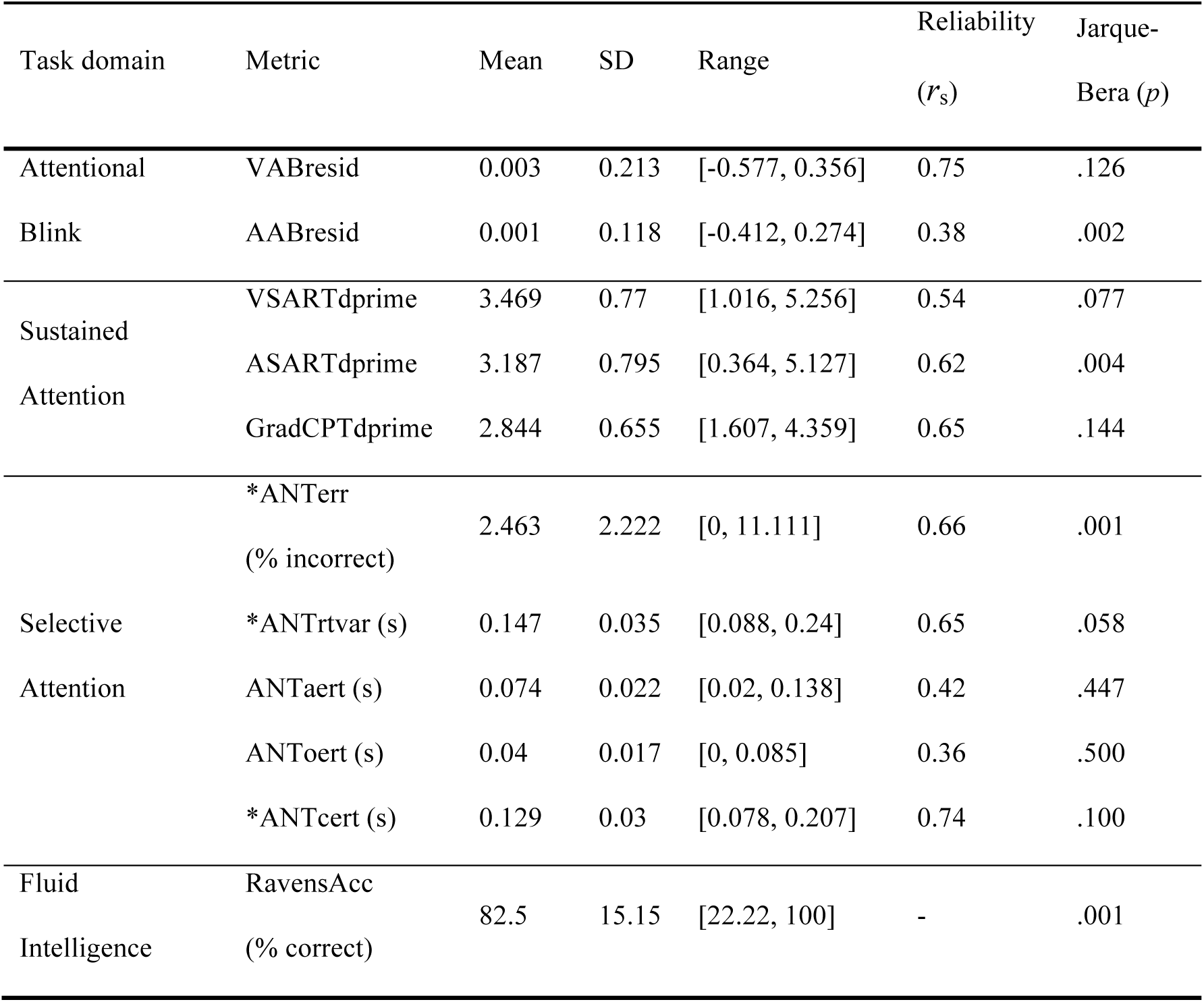
Summary of behavioral data. Test-retest reliability (Spearman correlations across sessions) was high for most metrics and significant for all (*p*s < .002). Jarque-Bera tests indicated some significant departures from normality. Metrics are presented before re-coding. Metrics that were subsequently reversed so that larger values would indicate better task performance are marked with an asterisk (*). N = 73 for all metrics except AABresid (N = 71).

To examine how behavioral performance was related across the various tasks, we computed Spearman’s rank correlations between each pair of behavioral scores (Figure 4). Significant correlations were primarily found within a given attentional domain. The three measures of Sustained Attention (VSARTdprime, ASARTdprime and GradCPTdprime) were significantly correlated, as were the primary measures of Selective Attention (ANTerr and ANTrtvar) and some of their component measures. Many of these metrics were significantly correlated, positively and negatively, across the Sustained and Selective Attention task domains as well. The negative correlations with the alerting (ANTaert) and orienting (ANToert) metrics may be due to their reflecting stimulus-driven attentional control, as opposed to the goal-directed control required for many other metrics. Attentional Blink (VABresid and AABresid) and Fluid Intelligence (RavensAcc) metrics generally were not significantly correlated with other metrics.

## Functional connectivity matrices

To assess and compare network connectivity during the VAB task and during rest, we plotted group-averaged functional connectivity (FC) matrices (Figure 5). The 419 parcellations from the FC matrices were matched to 17 network labels (Yeo et al., 2011), from which they were aggregated into eight cortical groups (Yeo, Tandi, et al., 2015) and a subcortical group. FC data from both the VAB task (VABFC; Figure 5a) and resting state (RSFC; Figure 5b) showed similar connectivity patterns, with largely positive within-network correlations and mixed directions for between-network correlations. The network correlation patterns were generally similar to those observed in other data sets (Yeo et al., 2011; Yeo, Tandi, et al., 2015). Using Network-based statistics to correct for multiple comparisons (Zalesky, Fornito, & Bullmore, 2010), we observed small differences between the two FC matrices, with many connections linking the Salience/Ventral attention and the Dorsal attention networks (Figure 5c). Such differences are consistent with the reported neural correlates of the attentional blink, which are in frontal and parietal areas associated with the ventral and dorsal attention networks (Marois, Chun, & Gore, 2000; Marois & Ivanoff, 2005).

**Figure 5.**
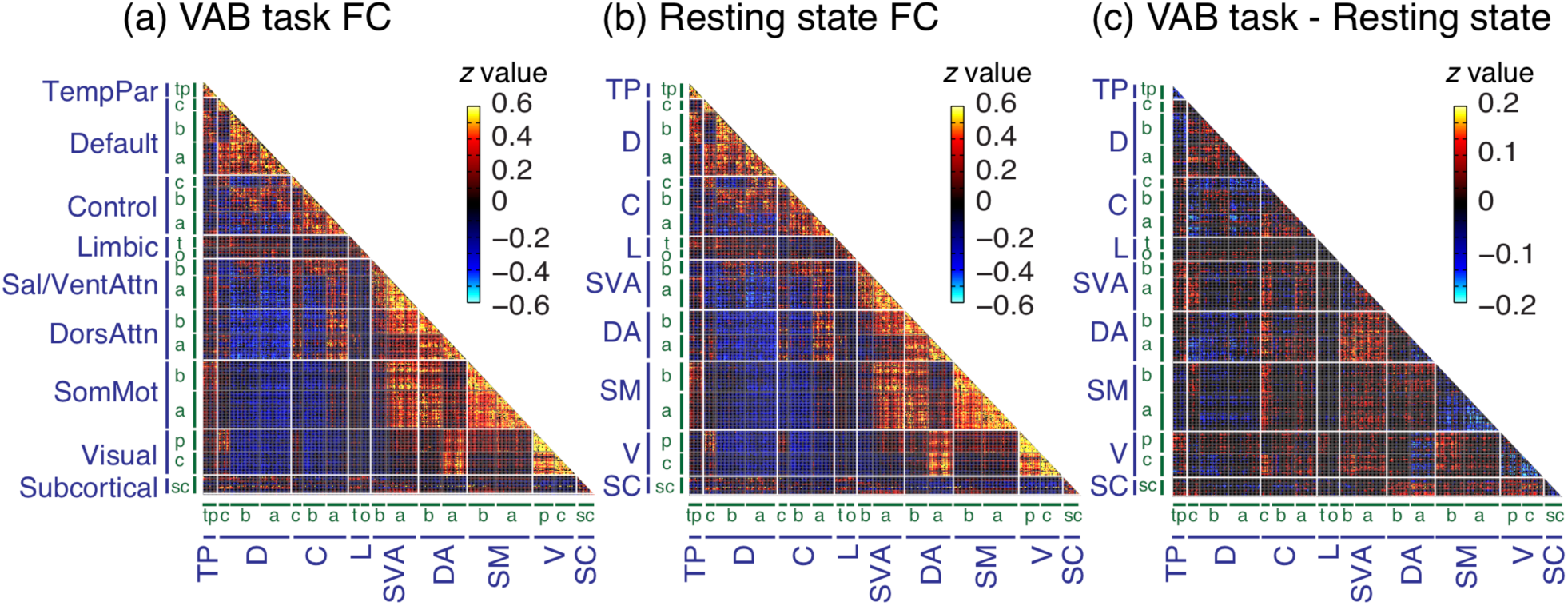
Functional connectivity (FC) matrices. Each edge was Fisher-transformed, and the resulting *z*-scores were averaged across participants. Edges found to correlate with motion were set to zero. (a) VAB task FC matrix (VABFC). (b) Resting-state FC matrix (RSFC). VABFC and RSFC patterns were similar to one another and to other data sets (Yeo et al., 2011; Yeo, Tandi, et al., 2015). (c) Difference FC matrix (VABFC - RSFC), showing edges that were significant at *p* = .05, corrected for multiple comparisons using network-based statistics. Differences between FC matrices were small, though they notably included connections linking the Salience/Ventral attention and the Dorsal attention networks. The 419 parcellations from the FC matrices were matched to 17 network labels (Yeo et al., 2011) (green labels), from which they were aggregated into eight cortical groups (Yeo, Tandi, et al., 2015) and a subcortical group (blue labels, spelled out in full in (a)). Subcortical regions include the brain stem, accumbens area, amygdala, caudate, cerebellum, ventral diencephalon, hippocampus, pallidum, putamen, and thalamus. For the green labels, letters represent the networks within the corresponding group, e.g., a(Default A), b(Default B), c(Default C), tp(TempPar), t(temporal pole in limbic region), o(orbital frontal cortex in limbic region), p(peripheral visual area), c(central visual area), and sc(subcortical).

## Behavioral predictions from CPMs constructed using the present dataset

Models constructed from VAB functional connectivity and behavioral data (vabCPMs) positively predicted VAB performance from task data (VABFC; Model type A) but not from resting state data (RSFC; Model type B) (Figure 6a; corresponding values are tabulated in Figure 6-1 in the Extended Data). Task predictions were made in relative performance ranks (Spearman correlations), so these predictions readily applied to other rank-order behavioral scores. When so applied, we found that Fluid Intelligence performance could be positively predicted from both task and rest FC data. Critically, the correlations between predicted and actual performance for Sustained Attention metrics were *negative*, as were these correlations for Sustained Attention tasks, albeit less consistently. Such results are counterintuitive, as all behavioral scores were re-coded so that larger values indicated better performance. Furthermore, one might expect that individuals whose network data predicted better performance on the VAB and Fluid Intelligence tasks would perform worse on Sustained Attention and Selective Attention tasks. In reality, however, VAB performance correlated insignificantly or weakly positively with other task metrics, whereas Fluid Intelligence performance was not significantly correlated with any other tasks. We return to this intriguing result in the Discussion.

**Figure 6.**
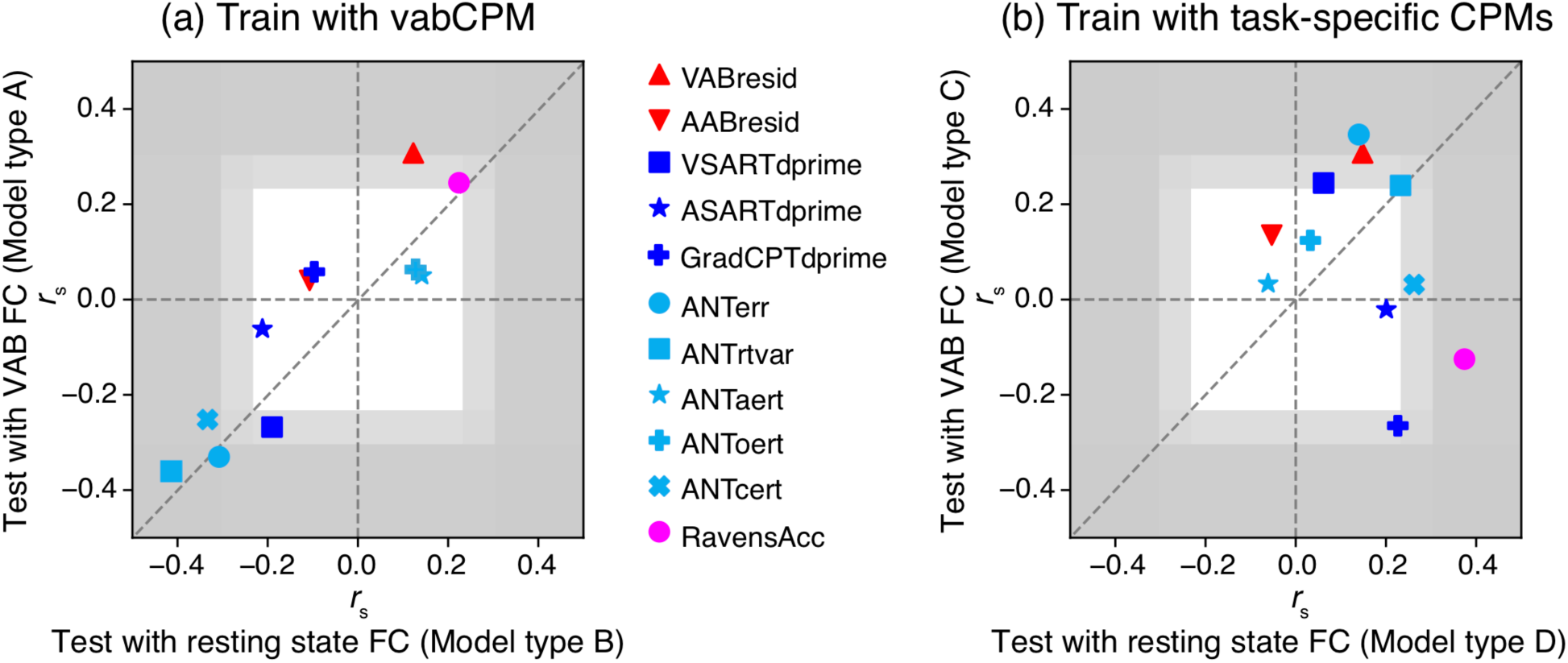
Behavioral predictions from CPMs. Each point represents a pair of Spearman’s rank correlation coefficients (*r*_s_) computed between observed and predicted behavioral scores for a given model type. (a) Predictions from vabCPMs, which were trained with VABFC and VABresid. Note the successful positive predictions for the VAB and Fluid Intelligence, but successful *negative* predictions for some Selective and Sustained Attention metrics. (b) Predictions from task-specific models. Many models could successfully predict behavioral performance, though results often varied greatly across the FC source. For both panels, the dark gray region indicates where *r*_s_ values are significant at the *p* = .01 level, and the lighter gray region indicates where *r*_s_ values are significant at the *p* = .05 level (uncorrected, with *d.f.* = 71). The *r*_s_ values and corresponding *p*-values are tabulated in Figure 6-1 in Extended Data. A standard edge selection threshold (*r*_threshold_ = .232, *p* = .05) was used for all models, though results were similar across a wide range of threshold values (Figure 6-2 and Figure 6-3 in Extended Data). Finally, as *p*-values from LOOCV procedures can be biased, we verified our results for the VAB using permutation testing; significance from this method and parametric approaches was consistent across edge selection thresholds (Figure 6-4 in in Extended Data).

Models built from task-specific behavioral data significantly predicted performance for each of the task metrics predicted from the vabCPMs, although the results were less consistent between VABFC-based (Model type C) and RSFC-based (Model type D) predictions (Figure 6b; corresponding values are tabulated in Figure 6-1 in the Extended Data). Although it is possible that the different predictions reflect different information in the two FC data sources (Figure 3), it is unclear whether the differences are stable or simply reflect difficulty in building models from fMRI data for which the behavioral data were collected separately. Indeed, CPMs constructed from fMRI data collected during task performance and that task’s behavioral scores tend to be more robust (Rosenberg, Finn, et al., 2016; Rosenberg et al., 2018; Yoo et al., 2017). Regardless, due to the same FC-behavior pair being used during both training and test, significant negative predictions from the vabCPMs became positive, as expected (e.g., VSARTdprime, ANTerr, ANTrtvat and ANTcert).

To examine whether model predictions were sensitive to the number of edges selected during CPM training, we explored how *r*_s_ changes as a function of edge selection thresholds. *R*_s_ values were observed to be reasonably stable across edge selection thresholds, though more variation was observed in task-specific models (Model types C and D) than in vabCPMs (Model types A and B) (Figure 6-2 and Figure 6-3 in Extended Data). Additionally, as *p*-values from LOOCV procedures can be biased, we verified our results for the VAB using permutation testing; significance from this method and parametric approaches was consistent across edge selection thresholds (Figure 6-4 in in Extended Data). Each of these issues may have contributed to the unexpected negative predictions for the GradCPTdprime metric, an aspect of the results to which we return below.

## Behavioral predictions from an external CPM (saCPM; Rosenberg et al., 2016)

For external validation of our model predictions, we applied the Sustained Attention CPM (saCPM) to our data. The saCPM was trained on fMRI and behavioral data from the GradCPT, a sustained attention task (Rosenberg, Finn, et al., 2016). When applied to our data, the saCPM predictions were virtually mirror images of our vabCPM predictions (Figure 7). Specifically, whereas the vabCPM predicted VAB and Fluid Intelligence performance positively and predicted Sustained Attention and Selective Attention performance negatively, the saCPM predicted VAB and Fluid Intelligence performance *negatively* and predicted Sustained Attention and Selective Attention performance *positively*.

**Figure 7.**
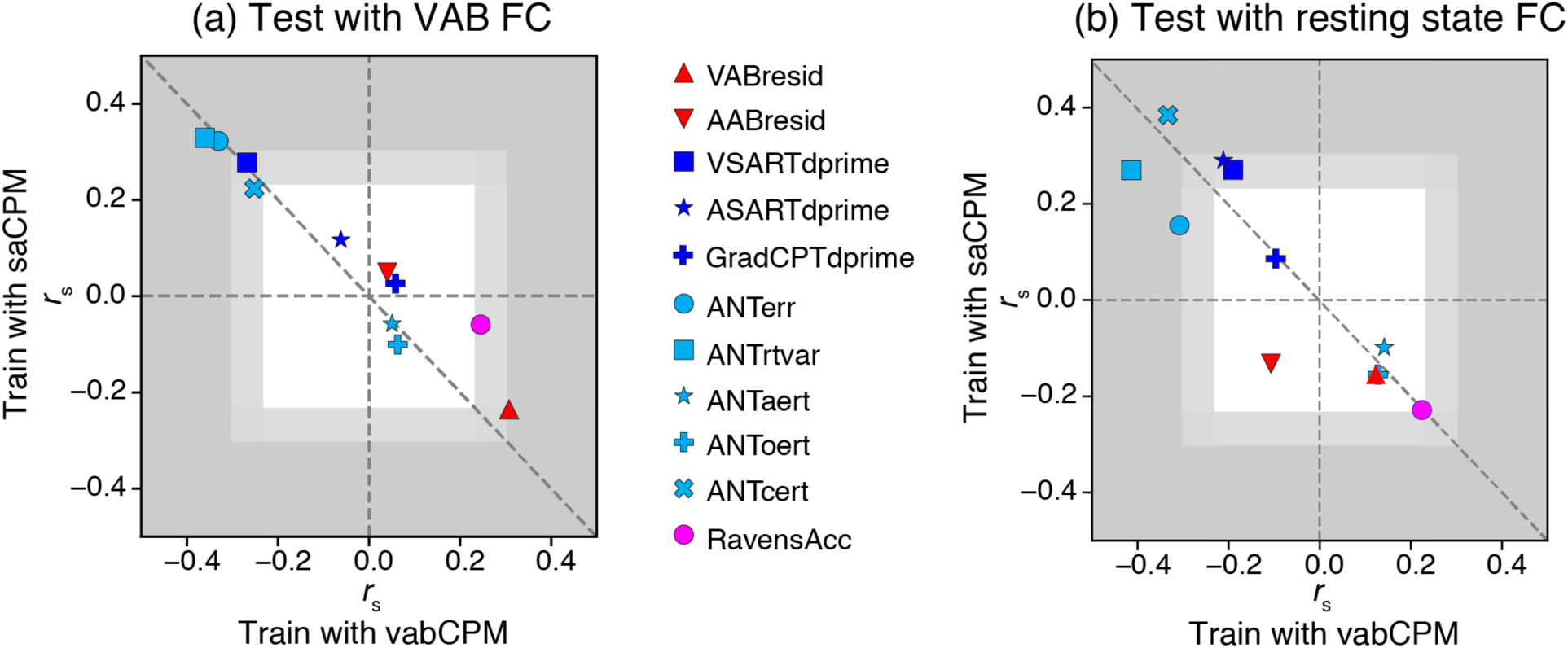
Comparison of predictions from the vabCPM and saCPM. Each point represents a pair of Spearman’s rank correlation coefficients (*r*_s_) computed between observed and predicted behavioral scores for a given model type. (a) Predictions from VABFC data. Note the saCPM’s successful positive predictions for some Selective and Sustained Attention metrics, with a successful *negative* prediction for the VAB. As noted above, the prediction directions were reversed for the vabCPM; indeed, the prediction points fall close to the diagonal. (b) Predictions from RSFC data. The vabCPM and saCPM predictions went in opposite directions, and were generally similar to the predictions from the VABFC. For both panels, the dark gray region indicates where *r*_s_ values are significant at the *p* = .01 level, and the lighter gray region indicates where *r*_s_ values are significant at the *p* = .05 level (uncorrected, with *d.f*. = 71). The edge selection threshold (*r*_threshold_) corresponded to *p* = .05 for all models. The *r*_s_ values and corresponding *p*-values for the saCPM are tabulated in Figure 7-1 in the Extended Data. (See Figure 6 for additional vabCPM details.)

With these findings, we also replicated the results from Rosenberg et al. (2018). Specifically, the saCPM was able to predict the error rates (ANTerr), reaction time variability (ANTrtvar), and conflict (ANTcert) metrics for novel individuals in the ANT task. Conversely, we failed to replicate the significant predictions for GradCPTdprime in our data, contrary to expectations. This replication failure is puzzling because the GradCPT behavioral data showed a reasonable spread of scores and good test-retest reliability (Table 3), and the saCPM and vabCPM did make significant predictions on other Sustained Attention metrics in our data set. We are currently exploring our GradCPT task and results further in our laboratory.

## Degree of overlap between predictive network pairs

To better understand how the underlying functional connectivity networks contributed to the model predictions, we analyzed the extent to which edges were shared between pairs of CPMs. For each pair, we calculated overlaps between positive network edges (those that predicted better behavioral performance for the model’s task), between negative network edges, and across positive and negative network edges (Figure 8). The networks derived from the vabCPM (VAB network) and saCPM (SA network) did not significantly overlap at similar network edges (e.g. positive-positive; Figure 8a and 8b). Instead, they significantly overlapped only at *opposing* network edges, the positive edges from one model and the negative edges from the other (Figure 8c and 8d).

**Figure 8.**
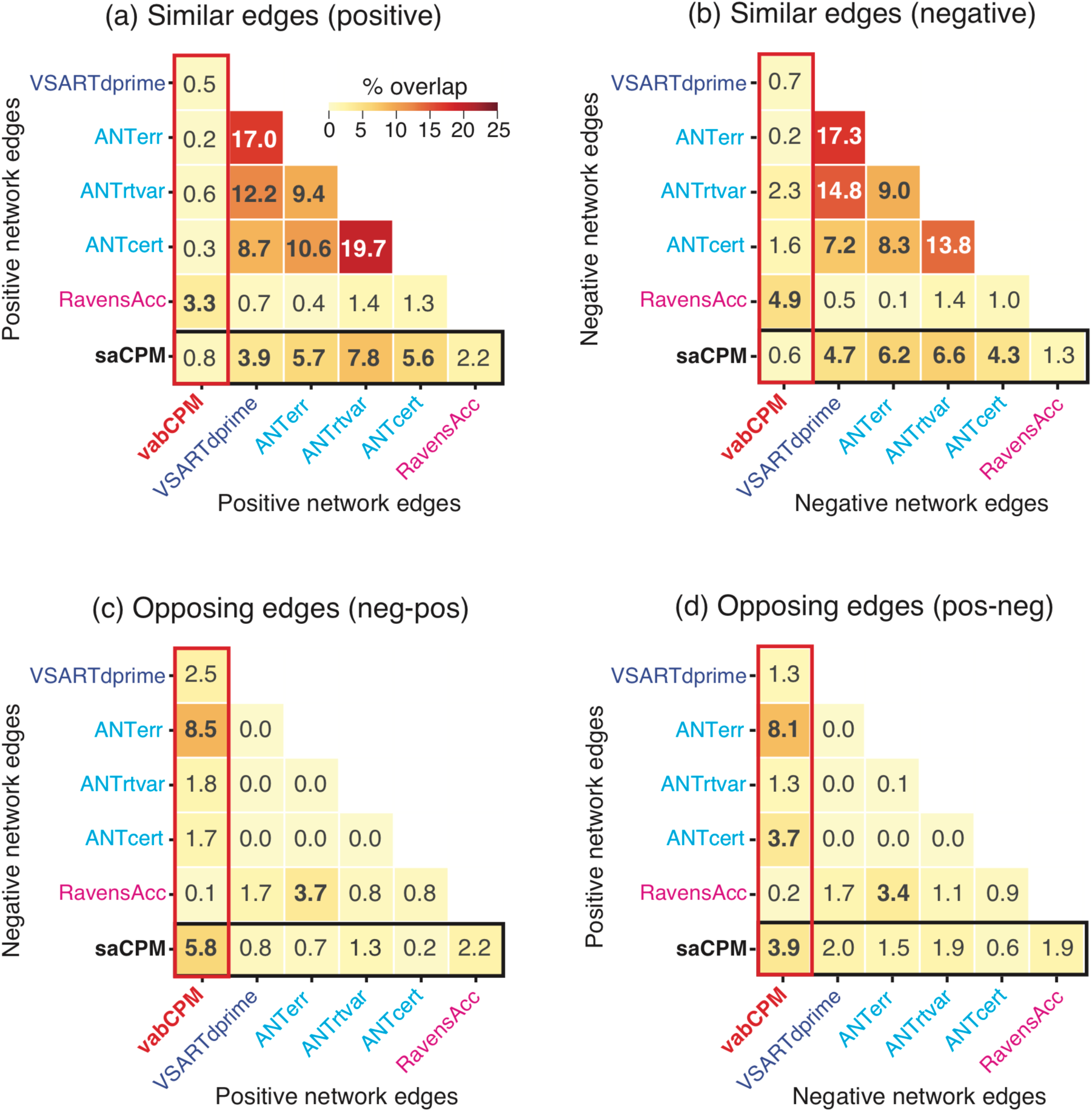
Percentage of edge overlap between networks from selected pairs of CPMs. Each task-specific model was based on VABFC data (Model type C). Model pairs with statistically significant overlap (*p* < .05, FDR corrected) are indicated in bold. Positive network edges predicted better behavioral performance for their associated metric, whereas negative network edges predicted worse behavioral performance. Overlap between *similar* network edges, (a) positive-positive and (b) negative-negative. Sustained Attention (including the saCPM) and Selective Attention models overlap primarily on similar edges, as do Fluid Intelligence and the vabCPM. Overlap between *opposing* network edges, (c) negative-positive and (d) positive-negative. The vabCPM network primarily overlaps with Sustained Attention (including the saCPM) and Selective Attention on their opposing edges. Only task metrics that were significantly predicted by vabCPMs using VABFC information are shown. The full set of overlaps for all CPMs can be found in the Extended Data (Figure 8-1, 8-2, 8-3, and 8-4). Note that the highest overlap value for any pairwise comparison, including each task metric predicted from VABFC and RSFC, was 26.5%.

The overlaps between the VAB network and SA network accorded with their overlaps with networks derived from other task-specific CPMs. VAB network edges tended to overlap more with the opposing network edges of the Sustained and Selective Attention models (Figure 8c and 8d) as compared to their similar network edges (Figure 8a and 8b). In contrast, SA network edges overlapped significantly with similar edges from each of the Sustained and Selective Attention models, but with none of their opposing edges. Finally, although VAB and Fluid Intelligence networks overlapped significantly only at similar network edges, SA network edges did not significantly overlap with either similar or opposing Fluid Intelligence network edges.

Taken together, the pattern of edge overlaps accords with the pattern of behavioral predictions. The VAB and SA networks overlapped at opposing edges, and their predictions were also negatively related (Figure 7). Predictions for individual metrics also aligned with model overlaps. In general, when a metric’s observed scores positively correlated with scores predicted from the vabCPM or saCPM, that metric’s CPM-derived network tended to overlap with the VAB or SA network at similar network edges.

We also observed significant overlaps between similar network edges from Sustained and Selective Attention models (Figure 8a and 8b). Similarly, Rosenberg and colleagues observed substantial overlaps between a high attention (positive) network from the GradCPT task and networks predicting high accuracy and low RT variability (better performance) in the ANT (Rosenberg et al., 2018). Taken together, these results suggest that sustained and selective attention share similar functional networks, at least in part. Such results also justify labels such as the “successful attention network” (Rosenberg et al., 2018). Importantly, these network overlaps are consistent with both the CPM predictions and the behavioral relationships (Figure 4). Significant network overlaps involving the VAB network, however, were not accompanied by significant behavioral relationships. We return to this observation in the Discussion.

## Anatomical locations of network edges

To better understand the networks that contribute to the successful behavioral predictions from the vabCPM and the saCPM, we investigated the anatomy of their underlying network edges. Briefly, for each pair of network groups (e.g. Visual and Salience / Ventral Attention), we expressed the number of shared connections (network edges) as a percentage of the number of possible connections between that pair (Figure 9). Edges that positively predicted VAB performance occurred primarily between the Default network and several other networks, including the Salience/Ventral attention, Dorsal attention, and Somatomotor networks (Figure 9a). For negative network edges, the most frequent connections occurred between the Salience/Ventral attention network and several other networks, including the Dorsal attention, Somatomotor, and Visual networks (Figure 9b). Note that such connections were also enhanced in VABFC compared to RSFC (Figure 5c), and that the connections across attention networks are consistent with the neural correlates of the attentional blink (Marois et al., 2000; Marois & Ivanoff, 2005). Negative network edges were also frequently found in Somatomotor network connections to the Dorsal attention network and to itself (within-network connections).

**Figure 9.**
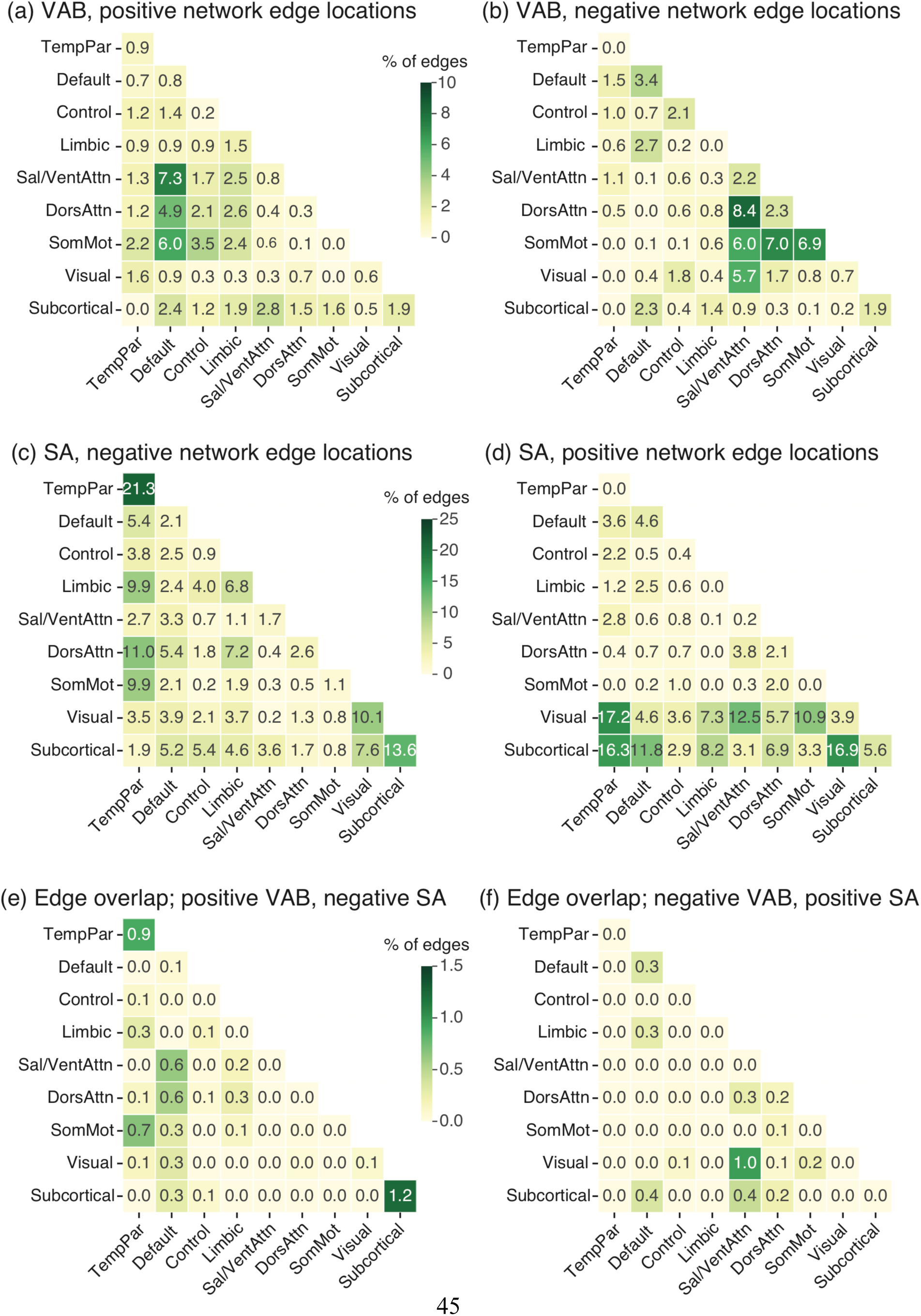
Anatomical locations of predictive attentional network edges. Each cell represents the number of shared connections between a pair of network groups, expressed as a percentage of the number of possible connections between that pair. For details about network groups, see Figure 5. (a) Positive and (b) negative VAB network edges. (c) Negative and (d) positive SA network edges. The panel order has been reversed for easier comparison with the opposing edges from the VAB network. (e) Positive VAB network edges and negative SA network edges (i.e. overlaps between (a) and (c)). (f) Negative VAB network edges and positive SA network edges (i.e. overlaps between (b) and (d)). Overlap percentages were numerically small, but included key network groups, particularly Default and Salience / Ventral Attention (Sal/VentAttn).

The SA network’s positive edges primarily included connections that involved the Visual and Subcortical network groups (Figure 9d). This pattern represents the remapping of the connections between the cerebellum and the occipital lobe of the “high attention” network (Rosenberg, Finn, et al., 2016) from the Shen-268 parcellation (Shen et al., 2013) to the Schaefer-419 one (Schaefer et al., 2017). Negative network edges included within-group connections in the TempPar and Subcortical networks (Figure 9c), which accord well with the intra-temporal, intra-cerebellar, and temporo-parietal connections in the “low attention” network (Rosenberg, Finn, et al., 2016). We observed additional negative edges within the Visual network and from the TempPar network to various other networks.

Although the VAB and SA networks involved largely dissociable sets of edges, some critical edges appeared to be shared (Figure 9e and 9f). Note that these overlaps were at opposing edges (Figure 8). Although overlap percentages were numerically small, they included key network groups. In particular, multiple identified connections involved the Default, Salience/Ventral attention, or TempPar networks.

## Discussion

### Predictive attentional networks

We used Connectome-based Predictive Modeling (CPM), a machine learning-based technique that associates task performance with functional connectivity measures, to construct a predictive model of Visual Attentional Blink (VAB) performance. Our model (vabCPM) successfully predicted VAB performance in novel individuals from fMRI data. The model’s predictions generalized to other task domains, including fluid intelligence (Finn et al., 2015).

Critically, vabCPM predictions for many sustained and selective attention task scores correlated *negatively* with the actual scores. As such, these significant predictions represent both model generalization and an extension of previous CPM results (Rosenberg, Finn, et al., 2016; Rosenberg et al., 2018; Yoo et al., 2017), but with new insights into attentional function owing to the divergent predictions.

For external validation of these results, we applied the Sustained Attention CPM (saCPM) (Rosenberg, Finn, et al., 2016), to our data. Previously, the saCPM successfully predicted task performance for sustained attention (GradCPT) (Rosenberg, Finn, et al., 2016) and selective attention (Attention Network Task, ANT) (Rosenberg et al., 2018). Here we broadly replicated these results: The saCPM successfully predicted sustained and selective attention task performance when applied to our participants’ data. In the sustained attention domain, however, significant predictions were found only for visual and auditory SART (Sustained Attention to Response Task) scores, but curiously not for GradCPT scores. In results that mirrored the vabCPM predictions, the saCPM predictions for VAB scores were negatively correlated with the observed scores.

This pattern of divergent predictions was also reflected in network overlaps. For the networks derived from the vabCPM and the saCPM, *opposing* network edges (i.e. positive from one, negative from the other) overlapped significantly, whereas similar network edges did not. Models constructed from each behavioral task and our fMRI data corroborated these results. As all behavioral data had been coded such that larger values indicated better performance, these divergent predictions indicate that “good” or “bad” network function was contingent on the task and its cognitive underpinnings. Moreover, the tasks were performed over several days, suggesting that the individual differences were stable and trait-like, not due to session-specific state effects.

## Implications for Cognitive Mechanisms

Although the observed pattern of CPM predictions may seem counterintuitive, it is consistent with both empirical evidence and theoretical positions. Foremost, our study’s attention tasks represent different ways of deploying voluntary attention. The VAB task requires rapid attentional engagement, disengagement, and re-engagement; sustained attention tasks require engagement over a prolonged period of time; the attention network task (ANT) requires the direction of attention to relevant spatial information. Skogsberg et al. (2015) proposed that the VAB and sustained attention tasks lie at opposite ends of a transient-sustained attention continuum. Rensink suggested that the VAB and ANT involve different core attentional functions: In the VAB, ‘attentional holding’ of one visual object leads to the failure to create a second visual object, whereas in the ANT, ‘attentional filtering’ selects spatial information (Rensink, 2013, 2015).

Empirical findings support the relative uniqueness of the VAB, while also suggesting that the ANT and sustained attention are more closely related. In our data, we found a general lack of behavioral correlations between the VAB and other attention task measures, consistent with previous studies of individual differences (Dale et al., 2013; Skogsberg et al., 2015). Conversely, we found moderately strong correlations between sustained attention tasks and the ANT. These tasks have been found to share similar functional networks (Rosenberg et al., 2018), a result we also replicated.

Nevertheless, the conceptual separation of the VAB from other attention tasks does not explain the *opposing* pattern of predictions from the same networks (saCPM and vabCPM), and the significant overlaps between their opposing network edges. Two observations provide important context for this finding. First, the VAB has spawned a variety of theoretical accounts (Dux & Marois, 2009), and its magnitude is sensitive to numerous disparate manipulations, ranging from requiring online responses (Jolicoeur, 1998) to concurrently listening to music (Olivers & Nieuwenhuis, 2005). As such, the VAB may have multiple causes. Second, our VAB predictions were generally moderate (*r* = .31 for vabCPM and *r* = .24 for saCPM; 5-10% of the variance), far smaller than the observed degree of stable individual differences (test-retest: *r* = .71; 50% of the variance). It is possible that the CPMs capture the variance associated with few, or even one, of the factors that affect VAB magnitude. If so, what could that factor be?

One plausible explanation is that our predictions reflect an individual’s propensity to maintain a more diffuse state of attention. This idea is consistent with the overinvestment hypothesis, in which the VAB results from too much attention being allocated to the first target; consequently, reducing attention on the RSVP stream improves performance (Dale & Arnell, 2010, 2015; Olivers & Nieuwenhuis, 2006). Similarly, task-concurrent mind-wandering, such as listening to music or thinking about a vacation, reduces the VAB deficit (Olivers & Nieuwenhuis, 2005, 2006). Similar effects are found in studies of disposition: Individuals with a greater propensity for mind-wandering tend to perform better in the VAB task (Thomson et al., 2015). Furthermore, mind-wandering has been linked to higher fluid intelligence and better problem-solving abilities (Baird, Smallwood, & Schooler, 2011; Godwin et al., 2017; Unsworth & McMillan, 2014), consistent with our CPM findings.

Diffuse attentional states, however, are associated with lower performance on tasks requiring more focused cognition. For example, in sustained attention tasks, individuals more prone to lapses in attention perform more poorly on the SART (Manly, 1999; Robertson et al., 1997; Smilek et al., 2010). Within individuals, distractive thoughts are associated with lower SART accuracy, prolonged and more variable RTs, and poorer response inhibition (Kam & Handy, 2014; Leszczynski et al., 2017; Stawarczyk, Majerus, Maj, Van der Linden, & D’Argembeau, 2011). In selective attention tasks, individuals more prone to mind-wandering performed worse on the ANT task (Gonçalves et al., 2017), and showed impaired exogeneous orienting of attention (Hu et al., 2012).

Both mind-wandering states and traits are also reflected in patterns of brain activity. Activity in the Default network and frontoparietal control regions increases during mind-wandering (Fox, Spreng, Ellamil, Andrews-Hanna, & Christoff, 2015). Similarly, individuals more prone to mind-wander had increased connectivity both within the Default network and between the Default network and frontoparietal control regions (Godwin et al., 2017). Such results are partially consistent with our findings. Edges between the Default and attentional networks (not the Control network) were related to positive VAB network edges and negative SA network edges.

On the weight of the available evidence, we propose that our vabCPM reflects individuals’ propensity towards diffuse attentional deployment. That propensity could indicate an individual’s ability to diffusely attend, their tendency to be in that state or mode, or both. We also suggest that other CPMs, including the saCPM, could reflect the complementary propensity towards more focused attentional deployment.

## Predictions from “Resting State”

Neuroimaging studies using ‘resting state’ data, in which subjects are scanned while not engaged in any particular task, have become increasingly popular. Initially used to identify functional architecture (Biswal, Yetkin, Haughton, & Hyde, 1995; Schaefer et al., 2017; Yeo et al., 2011), resting state data have recently been used to test whether functional architecture persists even when an individual is not engaged in a task that requires a given neurocognitive network (Finn et al., 2015; Jangraw et al., 2018; Lin et al., 2018; Rosenberg, Finn, et al., 2016; Yoo et al., 2017). Resting state studies also have many logistical advantages, including relatively easy standardization across multiple test sites and the potential for numerous applications from a single data set. In the current study, resting state CPMs (Model type D) replicated previous studies by successfully predicting sustained attention (Yoo et al., 2017), selective attention (Yoo et al., 2017) and fluid intelligence (Finn et al., 2015) task performance. We also replicated the result that models trained with *task-concurrent* FC data generally predict task performance better than models trained from resting state data (Yoo et al., 2017). Finally, when applying our trained vabCPM model to novel participants, predictions from task FC data were superior to predictions from resting-state FC data (Finn et al., 2017; Greene, Gao, Scheinost, & Constable, 2018; Rosenberg, Finn, et al., 2016; Rosenberg et al., 2018). This result might be due to amplification of behaviorally relevant individual differences in network patterns while performing a task (Greene et al., 2018).

## Methodological Considerations

Although we followed the guidelines in Shen et al. (2017) when developing our CPMs, there are two notable methodological differences. First, previous studies used the volumetric Shen-268 parcellation (Shen et al., 2013), but we used the surface-based Schaefer-419 parcellation (Schaefer et al., 2017). Second, previous studies have constructed linear models for associating network strength and behavioral scores, with predictions assessed using Spearman correlations (Fountain-Zaragoza et al., 2019; Lin et al., 2018; Rosenberg, Finn, et al., 2016; Rosenberg et al., 2018). We instead formed linear models from the ranks directly. Despite these differences, we still observed prediction patterns that were largely consistent and comparable to previous studies, providing evidence that CPMs are reasonably robust across such variations.

## Acknowledgements

The authors would like to thank Mike Esterman for the GradCPT experiment and analysis code. This work was supported by grants from the Singapore Ministry of Defence, DSO National Laboratories, and Singapore Ministry of Education and Yale-NUS College start-up, all to Asplund. It was also supported by an NUS Cross-Faculty Research Grant to Yeo and Asplund. The authors have no conflicts of interest, financial or otherwise, with respect to their authorship or the publication of this article.

## Extended Data

**Figure 6-1.**
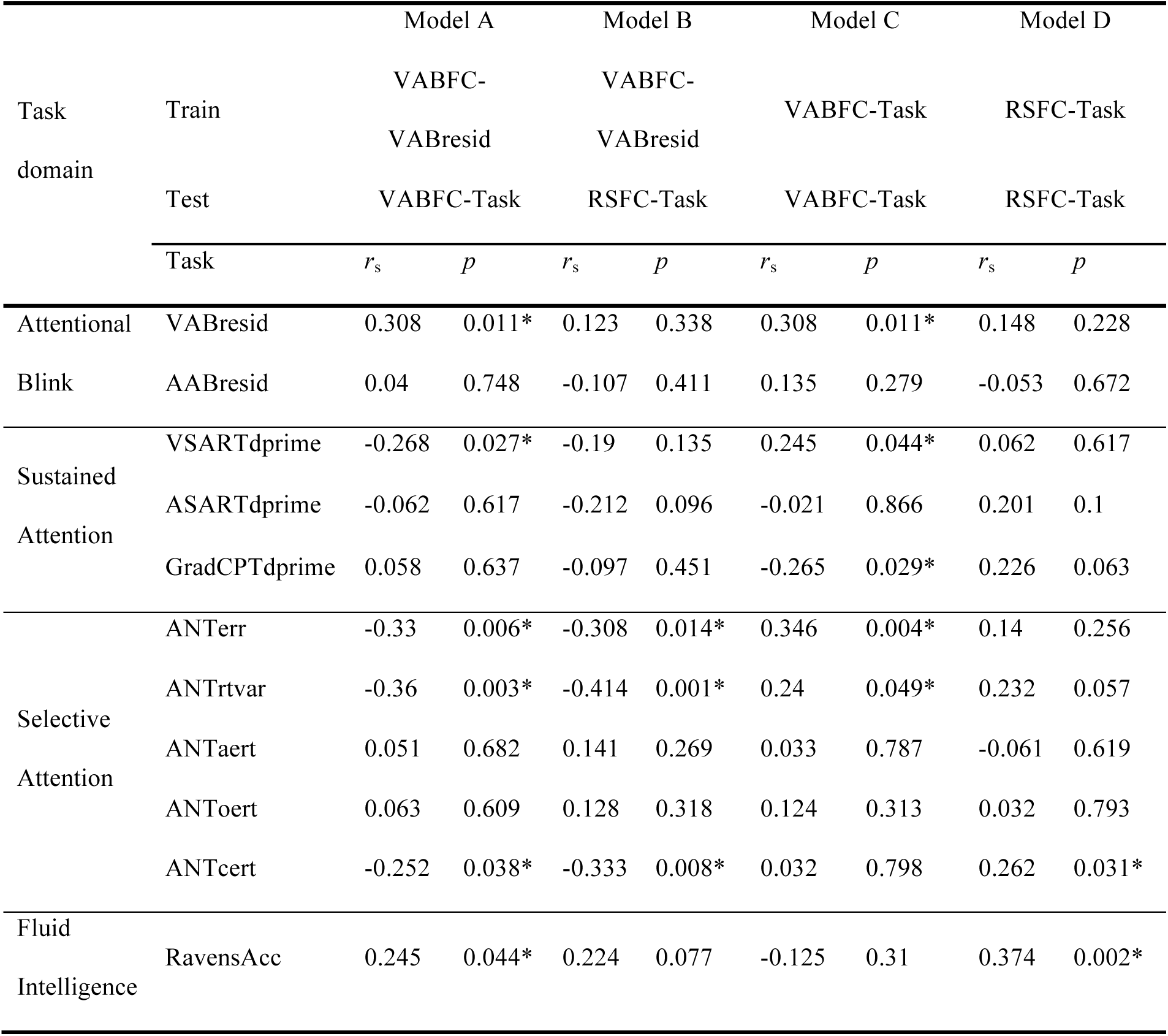
Behavioral predictions from CPMs. Values indicate *r*_s_ and uncorrected, two-tailed *p*-values from Spearman’s partial rank correlation, computed between predicted and observed behavioral scores, controlled for motion. The *p*-value corresponding to each *r*_s_ was found by transforming the correlation coefficients to Student’s *t* values by the partialcorr.m function in (*MATLAB*, 2014). Degrees of freedom for these tests were 66 for Models A and C, and 61 for Models B and D. (For AABresid, these *d*.*f*. values were 2 lower.) Note that all task scores had been re-oriented so that larger values indicate better task performance.

**Figure 6-2.**
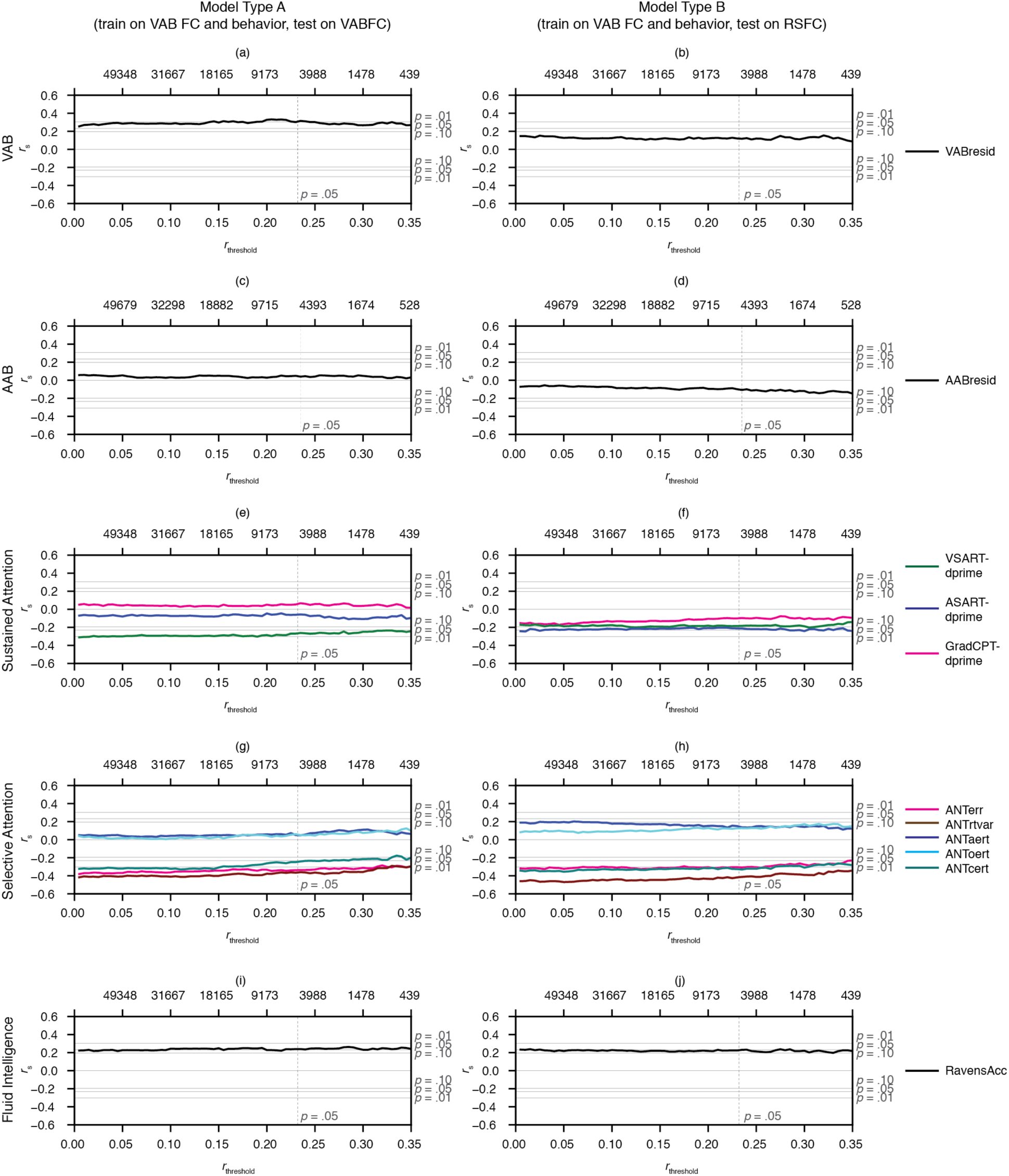
CPM predictions for models trained with VAB functional connectivity and behavioral data (Model types A and B) across edge selection thresholds. Within each subplot, the y-axis represents Spearman’s rank correlation values, *r*_s_, computed between predicted and observed task performance, controlled for motion. Horizontal gray lines indicate the corresponding *p* = .01, .05, .10, .10, .05, .01 (top to bottom) uncorrected levels of significance from standard *r*-to-*p* conversions, *d*.*f*. = 71 (69 for AAB). The x-axis represents edge selection thresholds, *r*_threshold_. The vertical gray line indicates the *r*_threshold_ at the *p* = .05 level of significance, *d*.*f*. = 70 (68 for AAB) due to one left-out participant during training. X-axis labels at the top of each plot indicate the average number of edges selected across all leave-one-out iterations at the corresponding *r*_threshold_ on the bottom x-axis.

**Figure 6-3.**
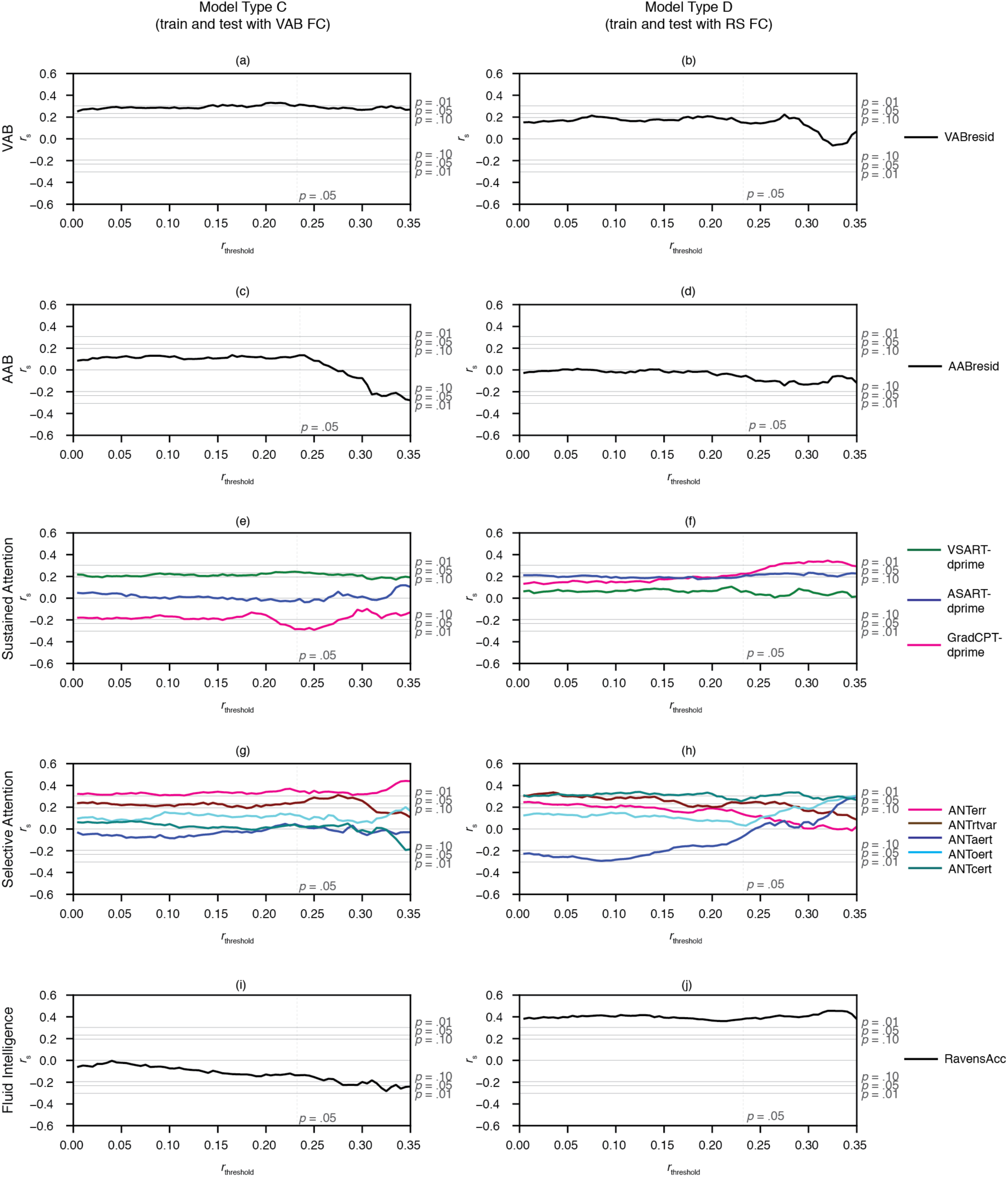
CPM predictions for models trained from task-specific behavioral data (Model types C and D) across edge selection thresholds. Within each subplot, the y-axis represents Spearman’s rank correlation values, *r*_s_, computed between predicted and observed task performance, controlled for motion. Horizontal gray lines indicate the corresponding *p* = .01, .05, .10, .10, .05,.01 (top to bottom) uncorrected levels of significance from standard *r*-to-*p* conversions, *d*.*f*. = 71 (69 for AAB). The x-axis represents edge selection thresholds, *r*_threshold_. The vertical gray line indicates the *r*_threshold_ at the *p* = .05 level of significance, *d*.*f*. = 70 (68 for AAB) due to one left-out participant during training. X-axis labels at the top of each plot indicate the average number of edges selected across all leave-one-out iterations at the corresponding *r*_threshold_ on the bottom x-axis.

**Figure 6-4.**
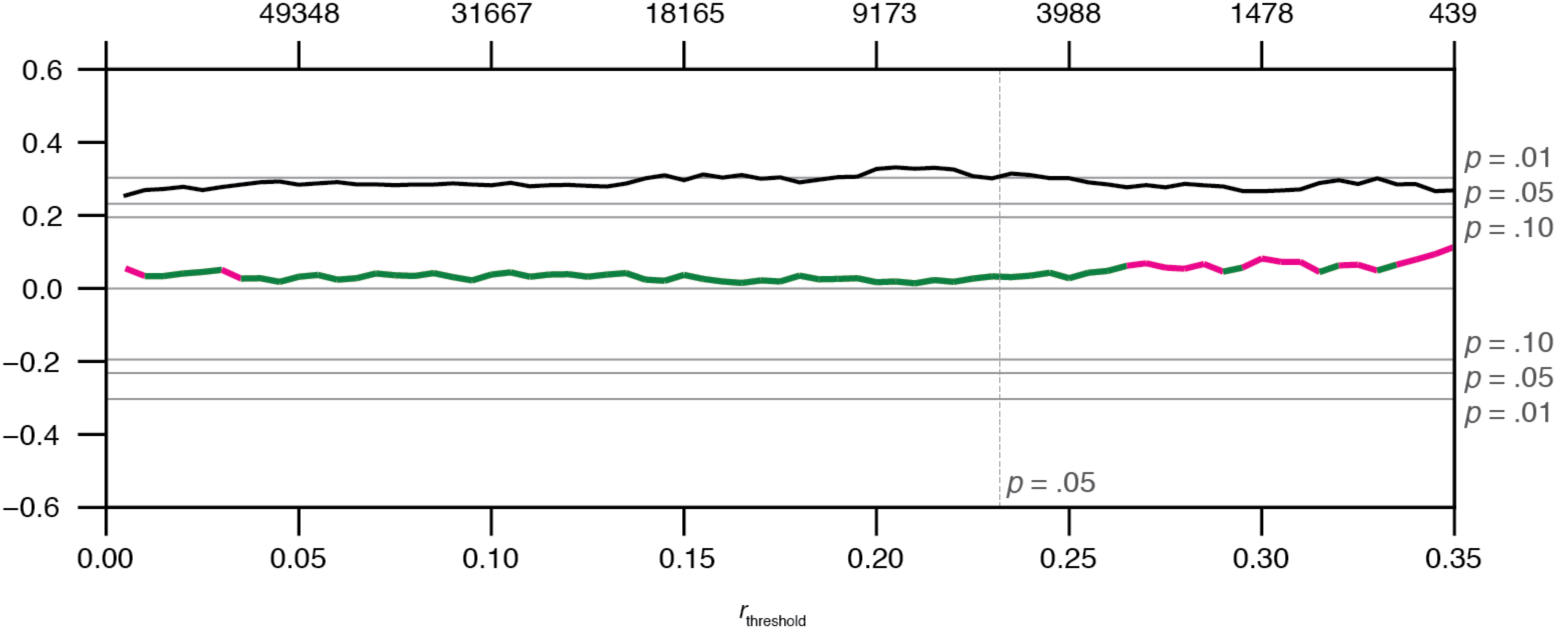
Permutation results of predicting VABresid data from VAB FC information, using the vabCPM (Model type A). The black line indicates Spearman’s rank correlation values (*r*_s_) computed between predicted and observed task performance, controlled for motion. Horizontal gray lines indicate the corresponding *p* = .01, .05, .10, .10, .05, .01 (top to bottom) uncorrected levels of significance from standard *r*-to-*p* conversions, based on *N* = 73. Green sections indicate *p* < .05 level of significance from permutation testing. Magenta sections indicate *p* ≥ .05 level of significance from permutation testing. Our analysis demonstrates a high level of consistency in the significance of *r*_s_ values between using standard *r*-to-*p* conversions (black line) and using permutation testing (green/magenta line) for the majority of edge selection thresholds.

**Figure 7-1.**
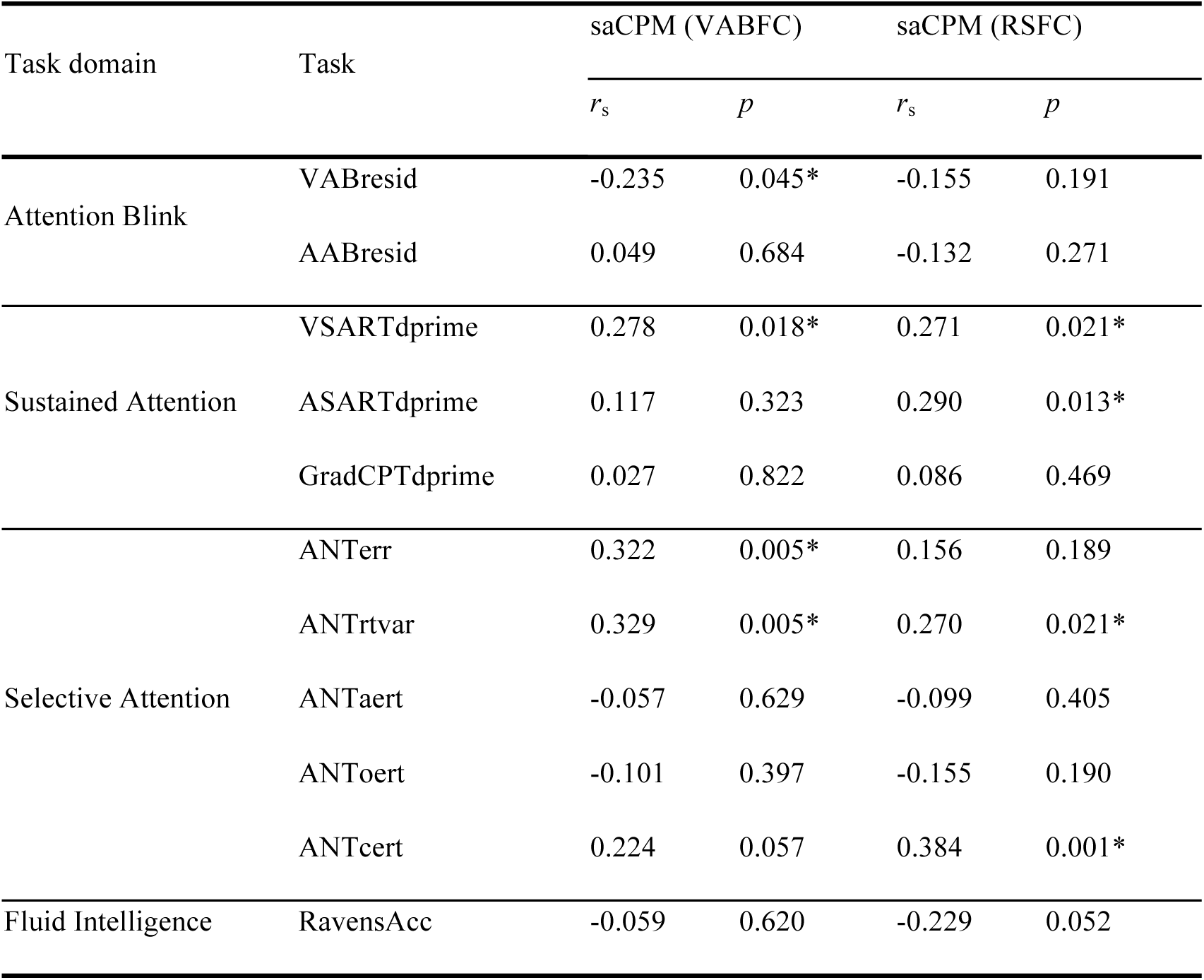
Results of applying the saCPM model to data in the present study. Values indicate *r*_s_ and uncorrected, two-tailed *p*-values from Spearman’s rank correlation, computed between predicted and observed behavioral scores. The *p*-value corresponding to each *r*_s_ was found using standard *r*-to-*p* conversions, with *d*.*f*. = 71 (69 for AABresid). All task scores had been re-coded so that larger values indicate better task performance.

**Figure 8-1.**
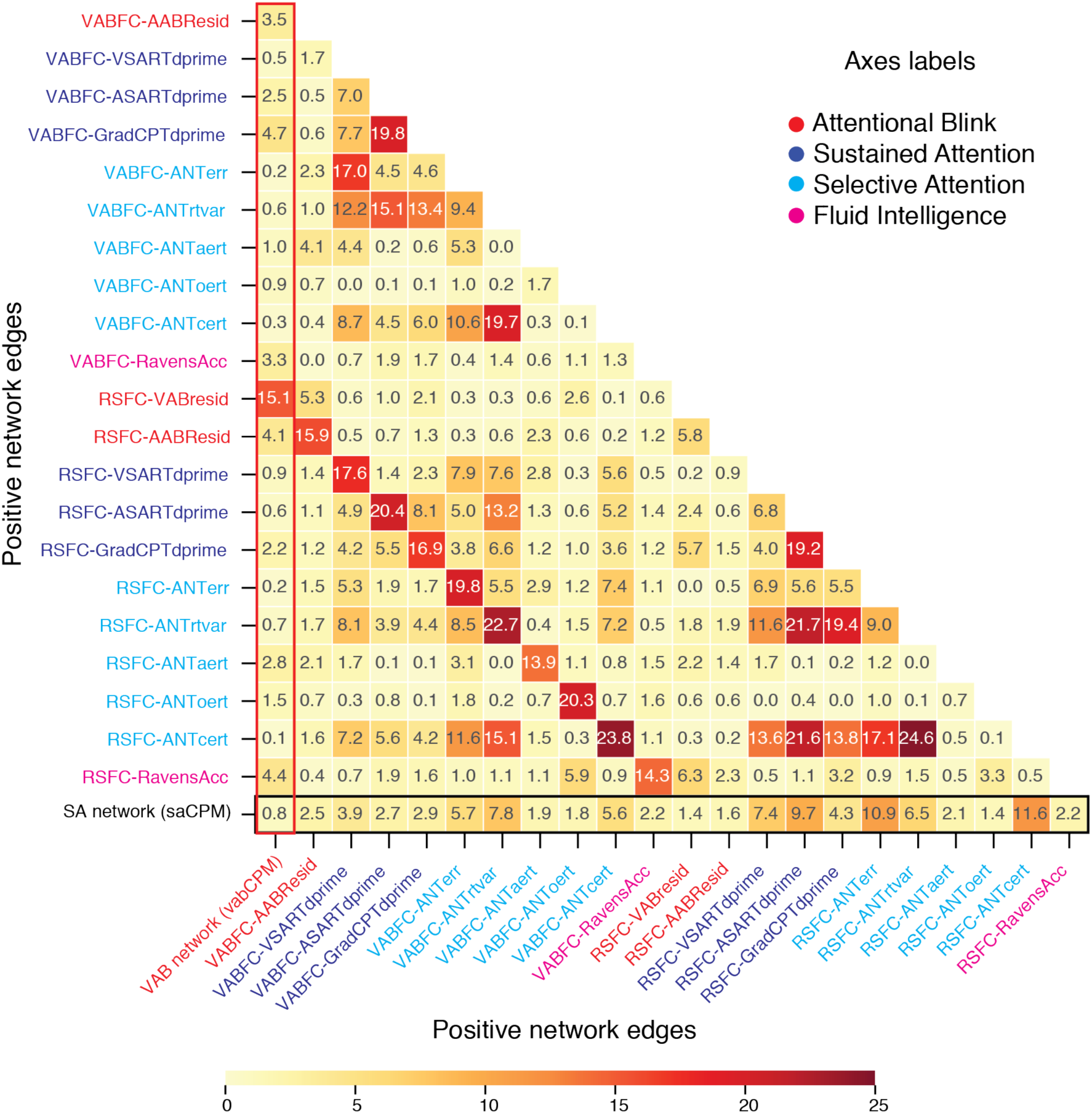
Percentage of edge overlap for positive network edges between each pair of models. The value within each cell indicates the percentage of overlap between the pair of models on the corresponding x and y axes. Overlaps between the vabCPM (saCPM) with the other models are illustrated in the column (row) bounded in the red (black) of each plot. Axes labels also reflect the FC-Task pair used for training the task-specific model.

**Figure 8-2.**
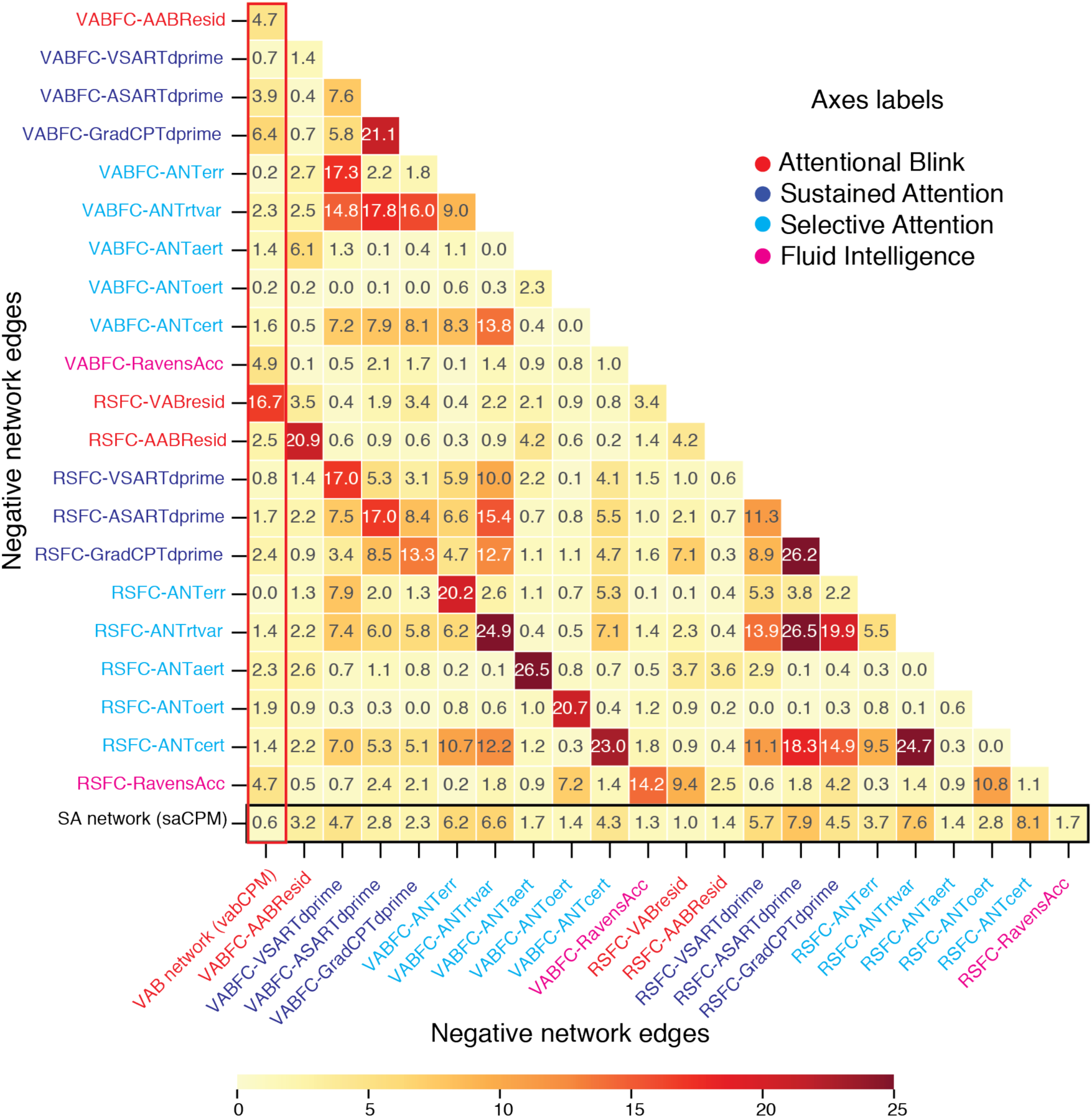
Percentage of edge overlap for negative network edges between each pair of models. The value within each cell indicates the percentage of overlap between the pair of models on the corresponding x and y axes. Overlaps between the vabCPM (saCPM) with the other models are illustrated in the column (row) bounded in the red (black) of each plot. Axes labels also reflect the FC-Task pair used for training the task-specific model.

**Figure 8-3.**
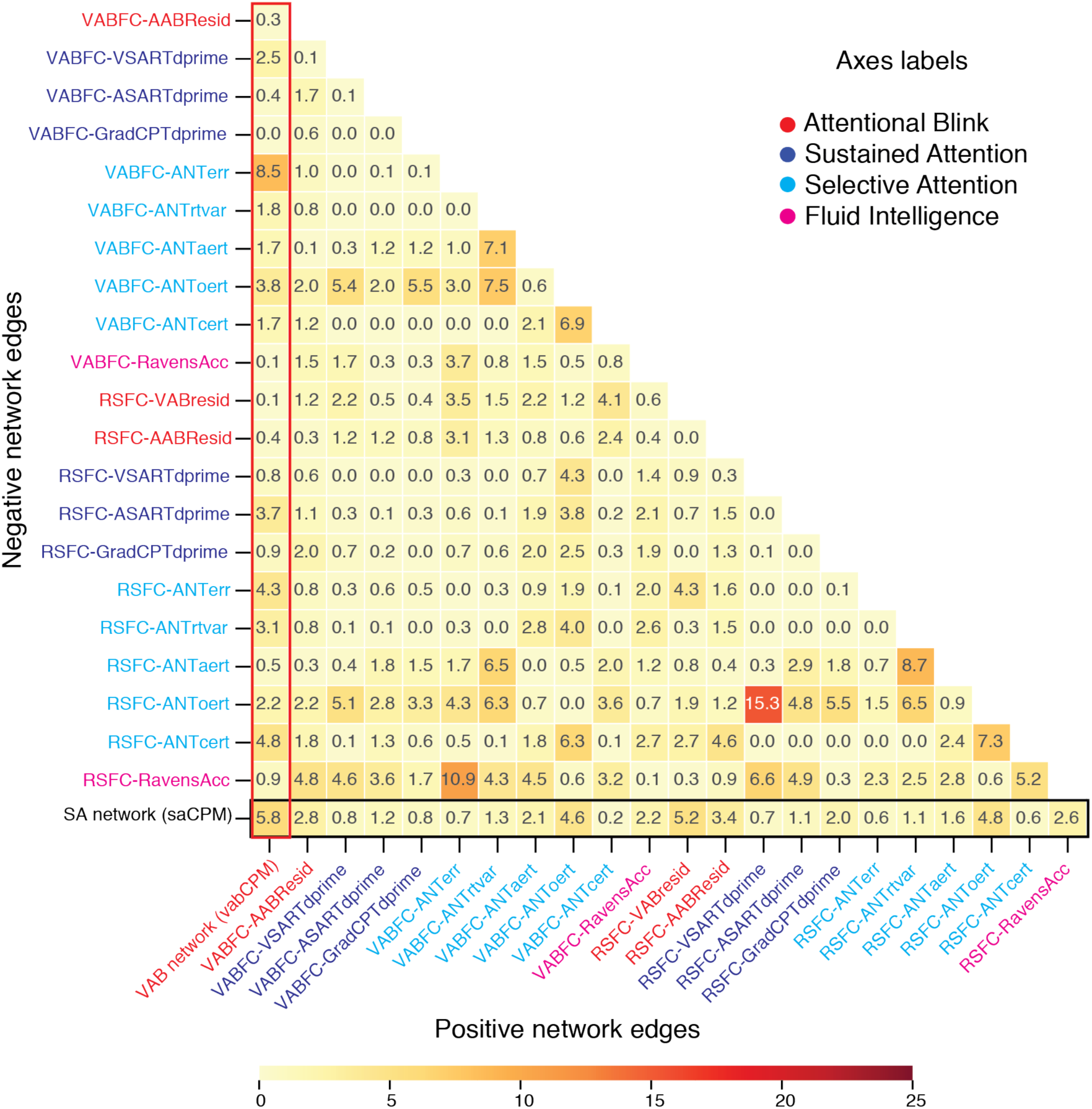
Percentage of edge overlap between negative network (y-axis) and positive network (x-axis) edges. The value within each cell indicates the percentage of overlap between the pair of models on the corresponding x and y axes. Overlaps between the vabCPM (saCPM) with the other models are illustrated in the column (row) bounded in the red (black) of each plot. Axes labels also reflect the FC-Task pair used for training the task-specific model.

**Figure 8-4.**
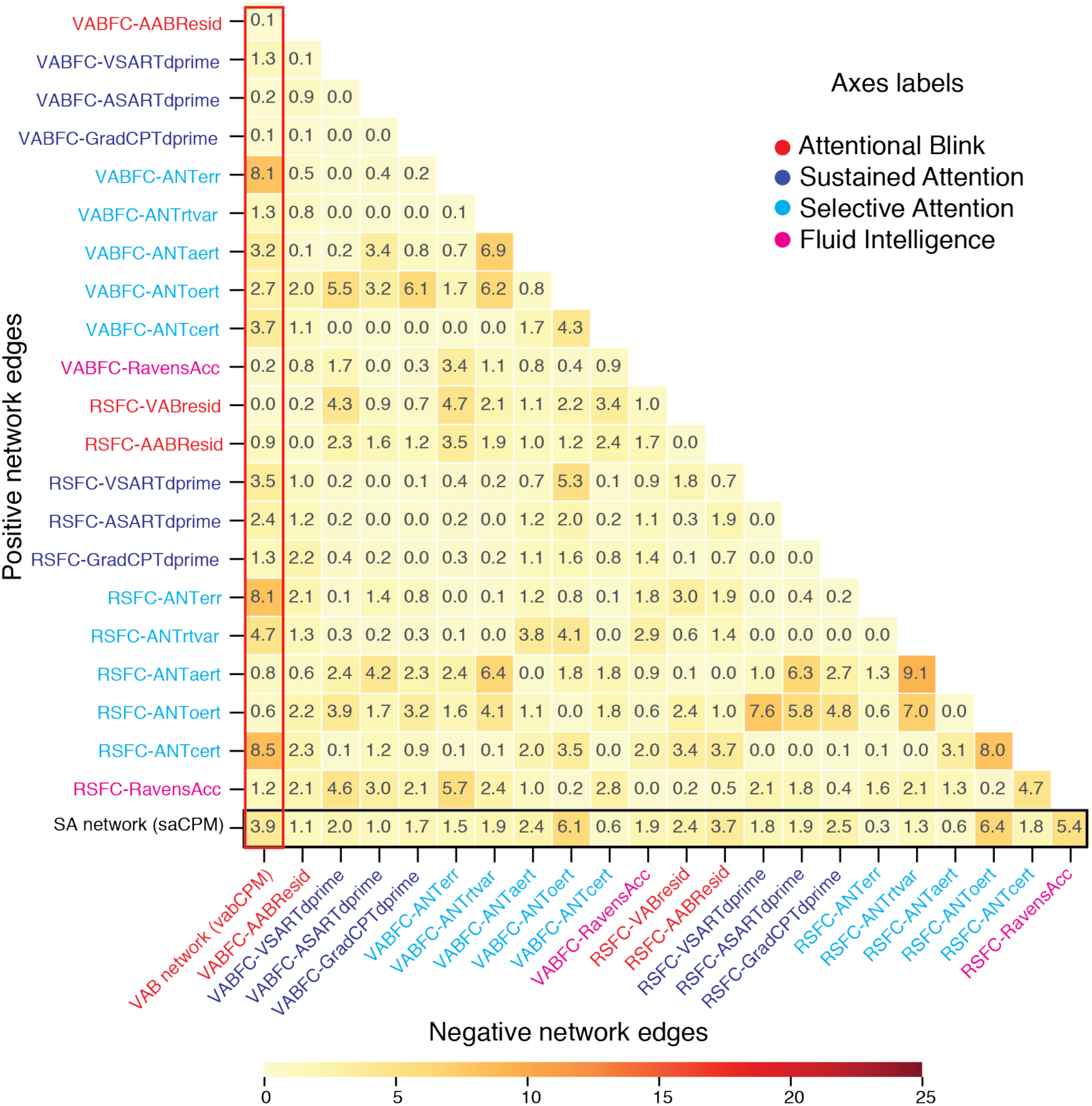
Percentage of edge overlap between positive network (y-axis) and negative network (x-axis) edges. The value within each cell indicates the percentage of overlap between the pair of models on the corresponding x and y axes. Overlaps between the vabCPM (saCPM) with the other models are illustrated in the column (row) bounded in the red (black) of each plot. Axes labels also reflect the FC-Task pair used for training the task-specific model.

**Table 1-1.**
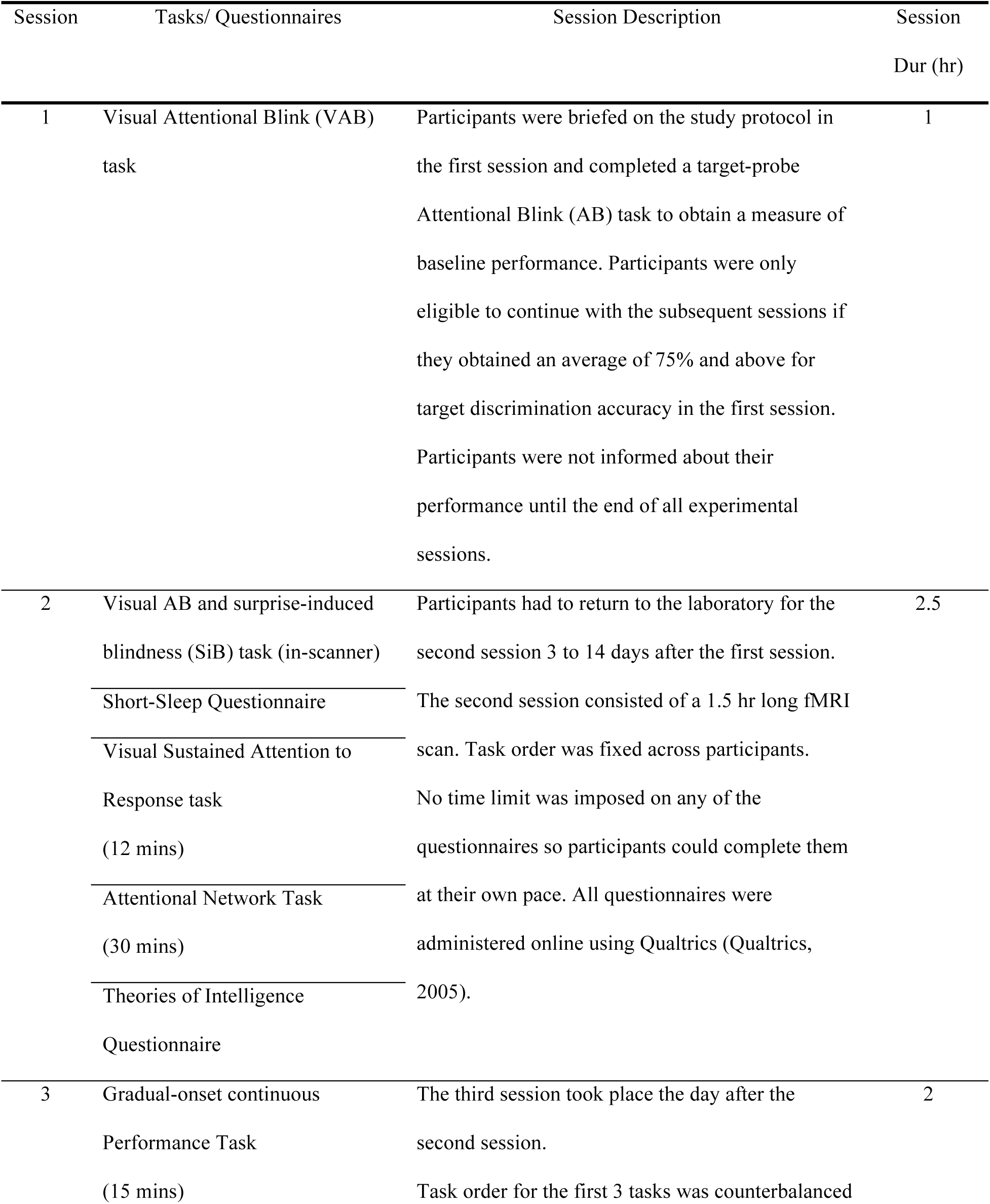
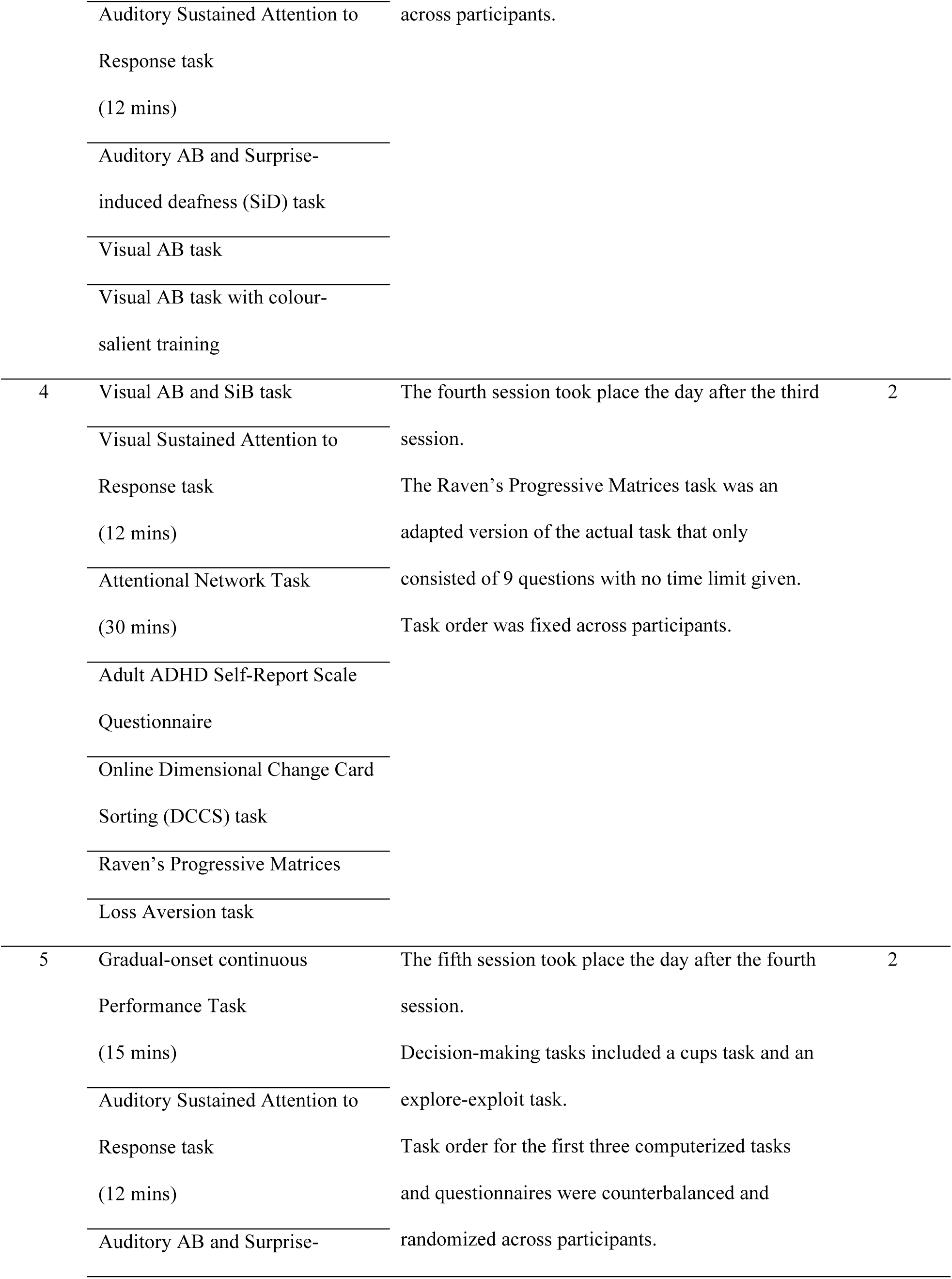
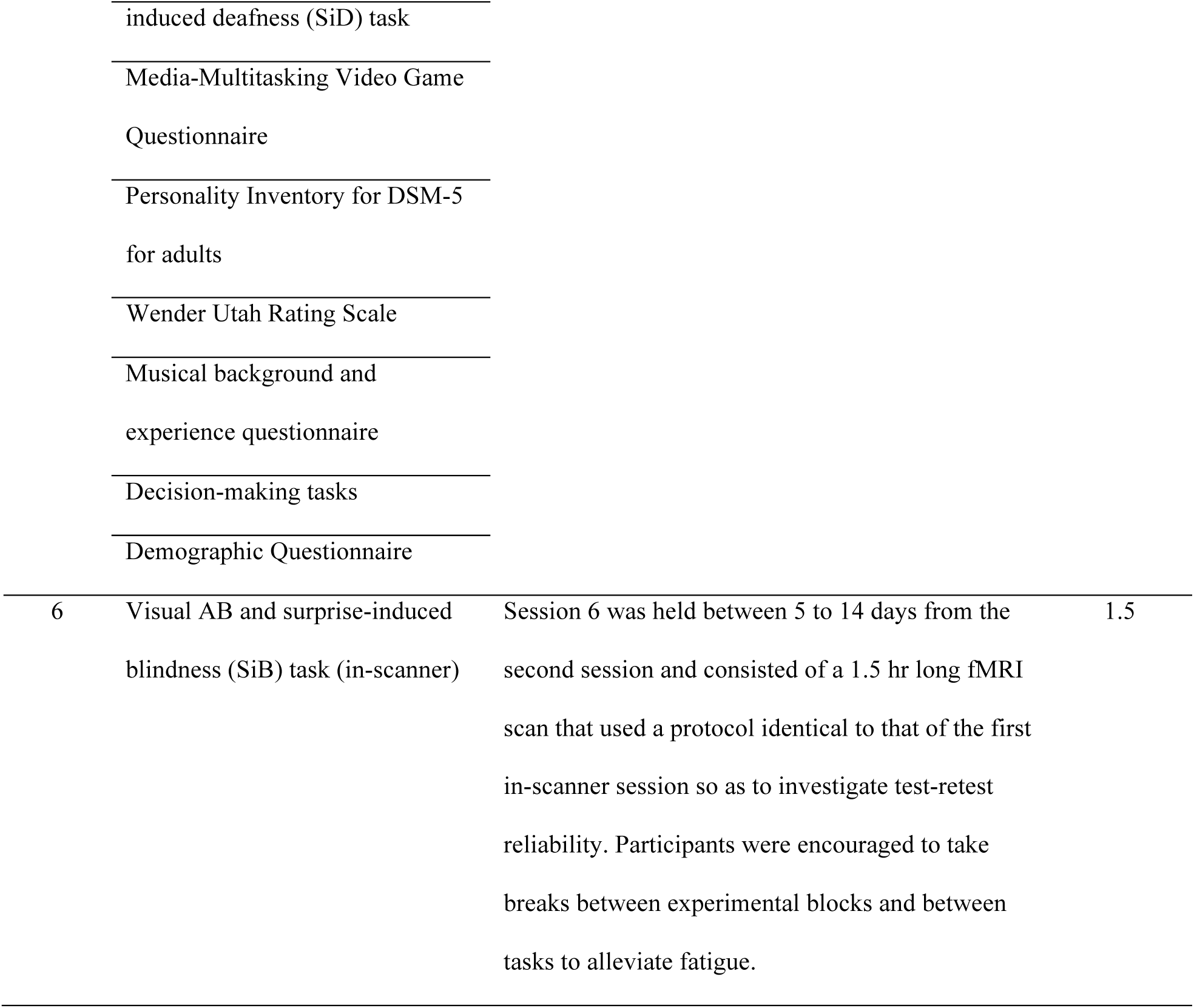
Detailed protocol from the full study from which the present data derives.

